# Acoustic Calibration in an Echoic Environment

**DOI:** 10.1101/321414

**Authors:** Alexander Kazakov, Nelken Israel

## Abstract

*Background*: The sound fed to a loudspeaker may significantly differ from that reaching the ear of the listener. The transformation from one to the other consists of spectral distortions with strong dependence on the relative locations of the speaker and the listener as well as on the geometry of the environment. With the increased importance of research in awake, freely-moving animals in large arenas, it becomes important to understand how animal location influences the corresponding spectral distortions.

*New Method*: We describe a full calibration pipeline that includes spatial sampling and estimation of the spectral distortions. We estimated the impulse responses of the environment using Golay complementary sequences.

Using those sequences, we also describe an acoustic 3D localization method for freely moving animals.

*Results*: In our arena, the impulse responses are dominated by a small number of strong reflections. We use this understanding to provide guidelines for designing the geometry of the environment as well as the presented sounds, in order to provide more uniform sound levels throughout the environment. Our 3D localization method achieves a 1 cm precision through the utilization of sound cues only.

*Comparison with Existing Methods*: To our knowledge, this is the first description of a large-scale acoustic calibration pipeline with acoustic localization for neuroscience studies.

*Conclusions*: Principled sampling of large arena allows for better design and control of the acoustic information provided to freely-moving animals.

## 1. INTRODUCTION

A growing number of studies in brain sciences use freely moving animals, a process that is driven by technological advances such as the possibility to perform extracellular recordings with large electrode arrays using telemetry^1,2^. However, less constrained behavior comes with a cost in terms of stimulus control. In audition, for example, sounds reproduced by loudspeaker are distorted as they propagate through the environment, so that the sound reaching the ear is different from that initially produced by the loudspeaker (Fig. 1a.). These distortions are usually estimated through acoustic calibration – the comparison between the sound at the input to the loudspeaker with that actually reaching the ears. For freely-moving animals, the need to calibrate sounds in multiple locations is an especially hard task.

Acoustic distortions are fully characterized by measuring the responses to an impulse (the ‘impulse response’). Evaluation of the acoustic properties using an impulse response is called ‘Impulse response analysis’, and its evolution is thoroughly described by Warren (2014)^3^. In short, the straightforward method to measure the impulse response is to record the responses to a single, ideal impulse. However, practical implementations of ideal impulses have low acoustic power because of their short duration, leading to low signal to noise ratio (SNR) in the resulting estimates. Instead of a single impulse, it is possible to use a pseudo-random noise, which is later cross-correlated with the recorded response, resulting in the impulse response of the system. For this to work well, the test signal has to have an autocorrelation consisting of a single peak at zero delay and zero correlation at all other delays. To maximize the quality of the estimates the test signal should also be of maximal power. For example, the use of maximal length sequences has been proposed by Schroeder (1979)^4^. Maximal length sequences, however, are not exactly white and thus can’t be used for estimating a perfect impulse response. To solve this problem, Golay complementary sequences were proposed^5^ and later applied for acoustic measurement in a number of studies of the auditory system^6–8^. Even though Golay complementary sequences require playback and recording of two different sequences, the test signal is shorter than its pseudo-random equivalents, resulting in faster estimation of the impulse response^5^. We therefore used Golay complementary sequences to achieve a high throughput calibration pipeline.

**Fig 1.**
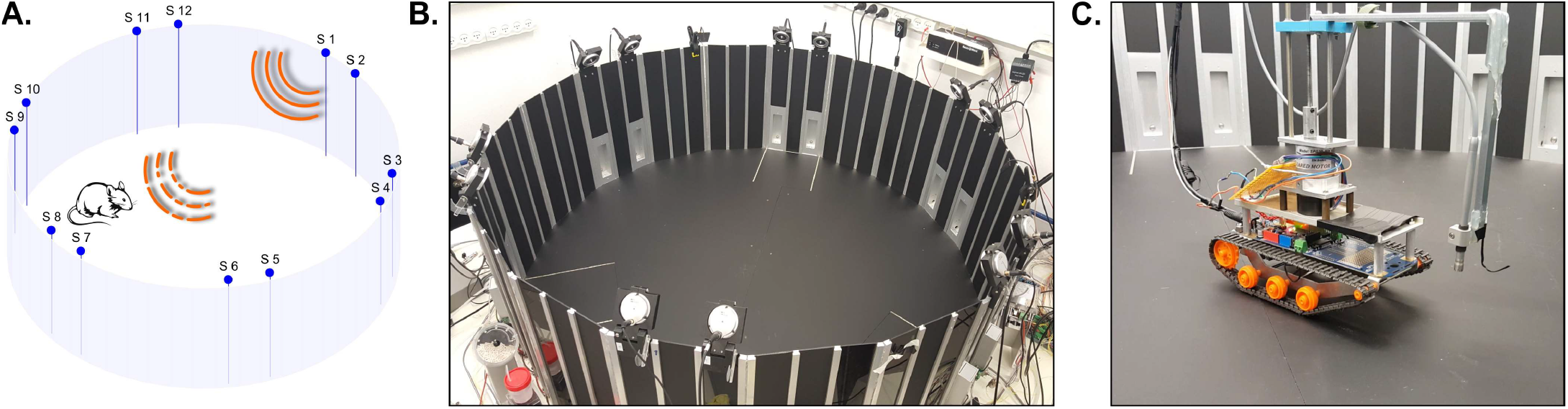
The echoic environment and the sound sampling robot. A. Simplified model of the arena, depicting the twelve speakers arranged in pairs (blue points along the circumferential wall). The arena is designed for experiments with freely behaving rats, driven by auditory stimuli (solid orange lines). Acoustic calibration compares the sound at the rat’s location to that transduced by the computer, and reflects both the speaker and the arena effects on the sound. B. An image of the arena. C. The acoustic calibration was performed in hundreds of locations along the arena, using a robot.

In this paper, we describe an automated pipeline for high throughput sound calibration in a large arena used in our own research. We demonstrate the use of acoustic calibration for understanding the geometrical origin of the resulting acoustic distortions, and illustrate some approaches to alleviate them. We then suggest a second use for the estimated impulse responses: we exploit them for rapid 3D localization of the animal. Fast and accurate localization of freely moving animals is an open challenge. Although many generic approaches have been proposed for object tracking, only few are applicable for studies of freely moving rodents. Tracking an animal in three dimensions (3D, volumetric tracking) is even harder. Camera based approaches require line of sight with the animal and installation of multiple cameras^9,10^. Solutions based on magnetic tracking require installation of dedicated sensors and may not provide sufficient spatial resolution^1^. Here, the same hardware used for acoustic calibration (multiple loudspeakers with a microphone on the head of the animal) is used also for high-resolution, fast 3D tracking.

## 2. MATERIALS AND METHODS

### 2.1 The arena

The arena is roughly circular (the circle is approximated by 18 straight segments, with diameter of 160 cm and wall height of 50 cm, see Fig. 1a-b). Six pairs of speakers (MF1, TDT) are evenly spread around the upper rim of the wall, with distance of 17 cm between speakers of each pair and central angle spread of 60º between successive pairs of speaker. All speakers are tilted by 35º below the horizontal axis such that their acoustic axes are pointed towards the center of the arena (Supp. Fig. 1).

### 2.2 Acoustic modeling of the arena

We idealize the acoustics of the arena as a linear, time-invariant system. The system is therefore fully specified by the impulse responses (IRs) from each speaker to each point in the arena. While the linearity assumption is very good at the sound levels considered here, the time invariance is clearly an approximation, since the presence of an animal in the arena modifies the acoustic properties in a way that depends on animal location. Nevertheless, the structure of the impulse responses is primarily determined by geometric factors that are independent of animal location. This point is further addressed in the Results.

The main challenge here is the need to sample the impulse response from multiple speakers to densely spaced points in the arena. Full reconstruction of the pressure field at all relevant frequencies requires the distance between spatially sampled points to be a fraction of the wavelength at the highest frequency of interest. However, this is practically impossible: rats hear up to 75 kHz, where the wavelength is 4 mm, requiring a sampling density of about 1 mm^11^. In practice, we sampled the IRs with spatial resolution of about 1 cm, using a custom built robot (Fig. 1c). While this resolution limits the ability to fully reconstruct the resulting pressure field to frequencies below 10 kHz, we will show that useful information can nevertheless be extracted at higher frequencies as well.

The Fourier transform of the IR (Fig. 2a) is called the transfer function (TF, Fig. 2b). The transfer function describes the steady-state amplitude gain and phase shift of sinusoidal signals as a function of frequency. Here we were primarily interested in spectral distortions, and therefore TFs are invariably represented by their amplitudes only. The TFs varied with the relative location of the microphone and the speaker that produced the IRs. We demonstrated this both by repositioning the microphone inside the arena, while estimating the IRs from a single speaker (Fig. 2c), or by measuring the IRs from different speakers at a microphone in a fixed location (Fig. 2d). The frequency-dependent variations are not produced by the noise in the measurement, since these were highly reproducible for a fixed location (Supp. Fig. 2).

### 2.3 Impulse Response measurements using Golay sequences

The IRs were measured by playing two complementary Golay sequences, as suggested by Zhou (1992)^6^ and illustrated in Supp. Fig. 3. Starting from *a*_1_ = [1 1] and *b*_1_ = [1 − 1], Golay sequences of length 2^*i*+1^ were constructed from Golay sequences of length 2^i^ following a recursive concatenation:

**Fig 2.**
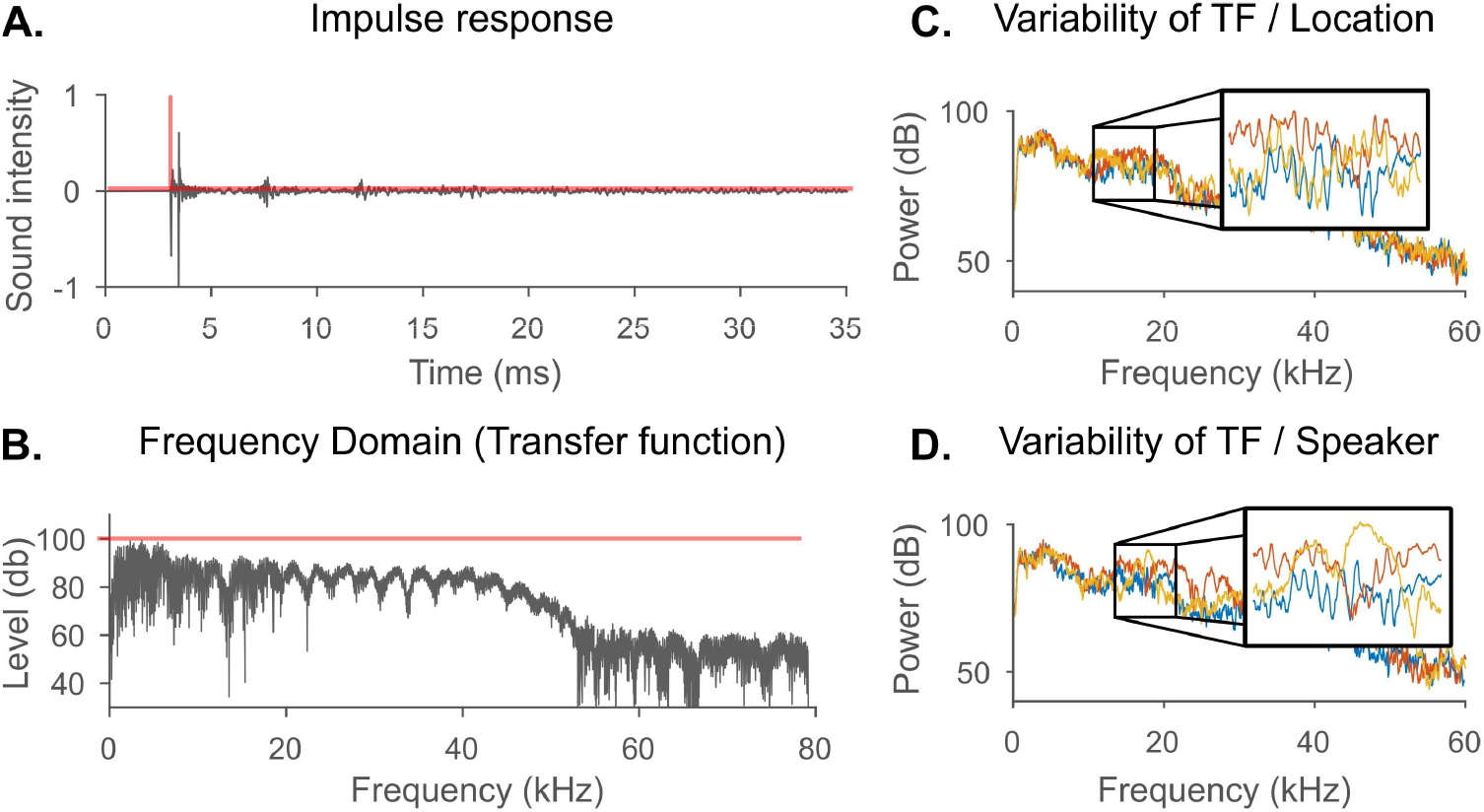
The acoustic properties of the arena were assessed by the impulse response analysis. The impulse response depends on the relative location of the microphone and the speaker that produces the impulse. **A**. An impulse response (IR) measured in the arena (black) and an ideal impulse response of a system with no distortions (red). **B**. The transfer function (TF) is the Fourier transform of the impulse response. Black and red lines show the TFs of IRs from A. **C**. Examples of TFs measured at 3 locations. **D**. Examples of TFs measured at a single location of the arena, for 3 different speakers.

*a*_*i*+1_=(*a*_*i*_|*b*_*i*_), *b*_*i*+1_=(*a*_*i*_|−*b*_*i*_) where (· | ·) denotes concatenation and (−) denotes elementwise negation.

These two sequences were played with a time interval of 0.2 s between them, to let most of the echoes decay (this interval is referred to below as the ‘silence buffer’). The sounds were generated by a single multichannel sound card (RME M16AD) and fed through programmable attenuators (PA5, TDT). Sounds were presented through TDT MF1 speakers. These speakers have a nominal frequency response range of 1–65 kHz with differences of up to ± 13 dB in sound level^12^. The speakers were driven by power amplifiers (SA1, TDT). Except for online tracking of rats, the sound was recorded by a calibrated microphone (Brüel & Kjær model 4939) at 192 kHz. The IR was recovered by cross-correlating the recorded sounds with the played sequences and summing the results obtained for the two sequences.

To ensure precise temporal synchronization of the playback and the microphone recordings, a square pulse was produced by the sound card on a separate channel and recorded on a separate channel of the sound recorder. Later this trigger was used to locate precisely the onset of the test sound, in order to estimate the length of the direct path of the sound from the loudspeaker to the microphone (Supp. Fig. 4).

### 2.4 Spatial sampling

Spatial sampling was automated using a custom-built robot, based on model RB-Rbo-33 by RobotShop (http://www.robotshop.com). The microphone was fixed to an extension arm in front of the robot, pointing perpendicularly towards the floor (Fig. 1c). A high-precision geared motor (Cytron Technologies, MO-SPG-30E-60K) was used to set the elevation of the microphone arm and was operated by an encoder (H bridge motor driver SN754410, Texas Instruments Inc.) connected to the Arduino board on the robot (Supp. Fig. 5). A custom MATLAB program controlled the automated sampling sequence: it positioned the robot in a predefined location and elevated the microphone to the desired height. It then played the test sound through each speaker in turn and recorded the microphone signals. The whole process repeated at the next location. As a rule, sampling at all microphone elevations occurred before moving the robot to a new location. In the data shown here, the spatial positions were selected in order to study particular features of the impulse response. For example, studying the floor echo was performed using 10 locations at varying heights at two horizontal locations (See results). When these measurements were performed, the robot was placed at the center of the arena and its arm was lowered such that the microphone was 1 cm above the floor. The arm was elevated 1 cm between each of the 10 vertical sampling locations, then the robot moved horizontally 40 centimeters (half the radius of the arena) and continued to sample the other 10 vertical locations, from the highest to the lowest one. The whole sequence was performed autonomously under control of a MATLAB routine.

The precision of the robot’s movements was evaluated and pre-calibrated to deliver sub millimeter accuracy per single movement. Nevertheless, positional error accumulated over a long measurement sequence, leading to potential errors on the order of 1 cm, and therefore limiting the length of recording trajectories. This error was partially eliminated, when needed, by breaking the recording session into a shorter sequence of autonomous sessions, with manual correction of the robot location between sessions. We didn’t integrate a camera feedback to correct the robot location since we expected its own error to be too large for that purpose (more than one cm).

### 2.5 Localization of a freely moving animal

The IRs were used to determine the time of arrival of the first waveform from each loudspeaker to the microphone, and transformed into an estimate of the distance between the microphone and the speaker (Supp. Fig. 4). The estimated distance constrains the speaker location to a sphere around the speaker. In order to narrow down the possible locations to a single point in space, distances to at least two other speakers are required (See Supp. Fig. 6; In general, if three spheres intersect, they would do so in 2 points, but in our case one point can be discarded since the microphone location is always below the plane defined by the speakers). The distance-based localization that relies on intersection of spheres is called trilateration (or multi-lateration for more than 3 distances). If the distances are exact, the relevant intersection point can be analytically found. In the presence of noise, the location of the microphone can be still numerically approximated. We used the MATLAB minimizer ‘fmincon’ with the loss function described by (1) to estimate the point *p*_*loc*_, where *p*_*i*_ is the location of speaker *i*, *d*_*i*_ are the calculated distances between the head stage and speaker *i*, and *p*^*^ is the estimated microphone location.

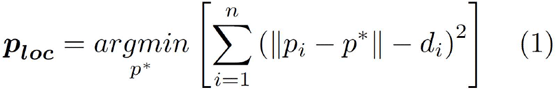

The animals used for localization were SABRA-R albino female rats. The animals used here had sufficient body mass (250–300 g) for carrying the headstage that included an integrated microphone and weighed 15 gram. The headstage was head fixed as described in Polterovich et. al. (In Press). The microphone was located 5 cm above the head, it sampled sound at 44 kHz and transmitted it online using a telemetry system to a data logging computer.

In addition to the acoustic localization, the arena was photographed at 30 Hz using a wide-field digital video camera that was placed above its center, such that each acoustic localization test could be later associated to a photograph of the rat in the RIFF taken at approximately the same time (see Supp. Movies 1–4). These images were later used to manually annotate the location of the microphone so that the localization error, defined as the distance between the position determined by acoustic localization and the manually annotated one, could be evaluated. The image of the rat was obtained through a fisheye lens, so that the height coordinate causes a shift outward from the vertical axis of the camera. In order to compare the manually annotated location of the headstage and the point estimated by the acoustic localization, the estimated location was shifted in the same way (Supp. Movies 1–4; The yellow point is the manual annotation of the microphone location, the red dot show the acoustically-determined position, corrected for the shift due to the elevation of the microphone. The red line displays the projection on the arena floor).

### 2.6 Additional Test Sounds

<u>Frequency sweeps.</u> Frequency sweeps covering 0.5 kHz extent in 10 seconds were used to directly measure the amplitude of the transfer functions over restricted frequency ranges. The ranges tested were 1–1.5 kHz, 10–10.5 kHz and 34–34.5 kHz.

<u>Narrow band sweeps.</u> Narrow band (NB) sweeps were created as a superposition of 101 pure tone sweeps. Each spanned the same range (0.5 kHz) but had a varying starting frequency and phase. For example, to produce NB sweep over the range of 10–10.5 kHz, we summed 101 frequency sweeps such that the first covered the range 9.95–10.45 kHz, the second covered 9.96–10.46 kHz, and so on, each starting with a random phase.

<u>Test sounds for spatial localization.</u> A variety of Golay codes of various lengths were tested for localization. The sequences were always sampled at 192 kHz. Since the microphone on the headstage could sample only at 44 kHz, the sequence was lowpass filtered at 20 kHz. To reduce measurement time, a shorter silent buffer of only 10 ms was inserted between the two Golay complementary codes. For a Golay code of length 2^12^ for example, this resulted in a total duration of sound presentation of 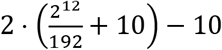 ms = 53 ms. Each speaker played these sounds and produced a single estimate of distance. Varying sets of 3 to 12 distances were used for estimating rat position. The total time for estimating three distances was 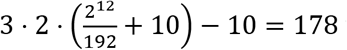 ms and for 12 distances (using all speakers) was 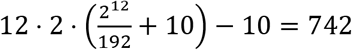 ms.

### 2.7 Data analysis

The signal processing and data analysis were performed using custom written MATLAB functions.

## 3. Results

### 3.1 Impulse response and transfer function encode local acoustic features

Perfect sound reproduction occurs only when the impulse response (IR) represents a pure delay; in that case the transfer function (TF) has an amplitude gain which is constant as a function of frequency, and a phase shift which is a linear function of frequency (Fig. 2a-b, red lines). Deviations from these ideal conditions cause the IR to deviate from a pure delay, and the TF typically has frequency-dependent amplitude gain (Fig. 2a-b, black lines).

One concern about these measurements is the potential role of the robot itself in shaping the IRs, since its body could produce its own share of acoustic distortions. To check this, we recorded IRs from the microphone at the same location while the robot rotated around it, showing only minor differences in the TFs and mostly at frequencies above 15 kHz: In a control experiment, the robot was rotated twelve times by 90º clockwise, keeping the microphone in the same position. When comparing the average IR produced by the speakers obscured by the robot (red plots, Supp. Fig. 7a.) with the IR of the speakers that had line of sight with the microphone (same figure, blue plots) it is evident that the crucial temporal features are preserved: The sound onset was unchanged (t = 3.1 ms), and the main features of the impulse response, the two amplitude deflections at t = 3.2 and t = 3.4 ms, are intact. The two TFs (Supp. Fig. 7b.) largely overlap up to 25 kHz. The recorded TFs from all locations shared many features (as can be seen on an illustrative TF in Fig. 2b): the TFs were invariably lowpass and the energy above 45 kHz was reduced by about 40 dB relative to low frequencies. This is most probably due to the frequency response of the loudspeakers^12^. Another feature is the slow oscillations with a period of a few kHz, easily observable between 20 and 40 kHz in Fig. 2b. In addition to the slow oscillations, there were fast fluctuations of the TF. For example, in the TF shown at Fig. 2b, sound level has a local minima near 13, 15 and 23 kHz, where the sound level decreased by 30 dB and then increased again within a frequency interval as narrow as 40 Hz, forming so-called spectral notches. Finally, TFs exhibited stronger attenuation of high frequencies when the microphone was located far from the acoustic axis of a loudspeaker.

We will show that the periodic patterns in the TFs (both slow and fast) are due to strong reflections from the floor and walls of the arena (Section 4.2), while the attenuation off the acoustic axis at high frequencies is the consequence of the frequency-dependent directivity of the speakers (Section 4.3). We then develop strategies to at least partially compensate for these distortions (Section 4.4).

### 3.2 Echoes from the floor and walls shape the IRs

The IRs tend to have a relatively simple structure, consisting of the direct sound reaching the microphone (indicated by red arrows in Fig. 3a) followed by a first prominent reflection (yellow arrow) and a few longer-latency reflections (the green and blue arrows mark the first two).

In order to verify the origin of each of these reflections, we converted the timing of arrival (ToA) of a reflection into traveling distance of the sound wave (distance = ToA × speed of sound at 20º C, considered as 343 m/s. For an example, see Supp. Fig. 4). The precision of the acoustically determined distances was tested in a separate control experiment, and the error was within the 0.3 cm uncertainty of manual measurements: The precision of the acoustically determined distances was tested by measuring the delay of the direct sound from all 12 speakers to the microphone at the center of the arena, comparing them to manually measured distances (Supp. Fig. 1 depicts geometric modeling of the arena). The measured ToA of the sound was 2.62 ms, corresponding to a traveling distance of 89.8 cm. The empirical distance was 90 cm, within less than 1% of the value determined from the ToA. Because of the speaker’s shape the empirical distance was determined with a spatial precision of 0.3 cm. All acoustically-determined distances fell within this error range.

**Fig 3.**
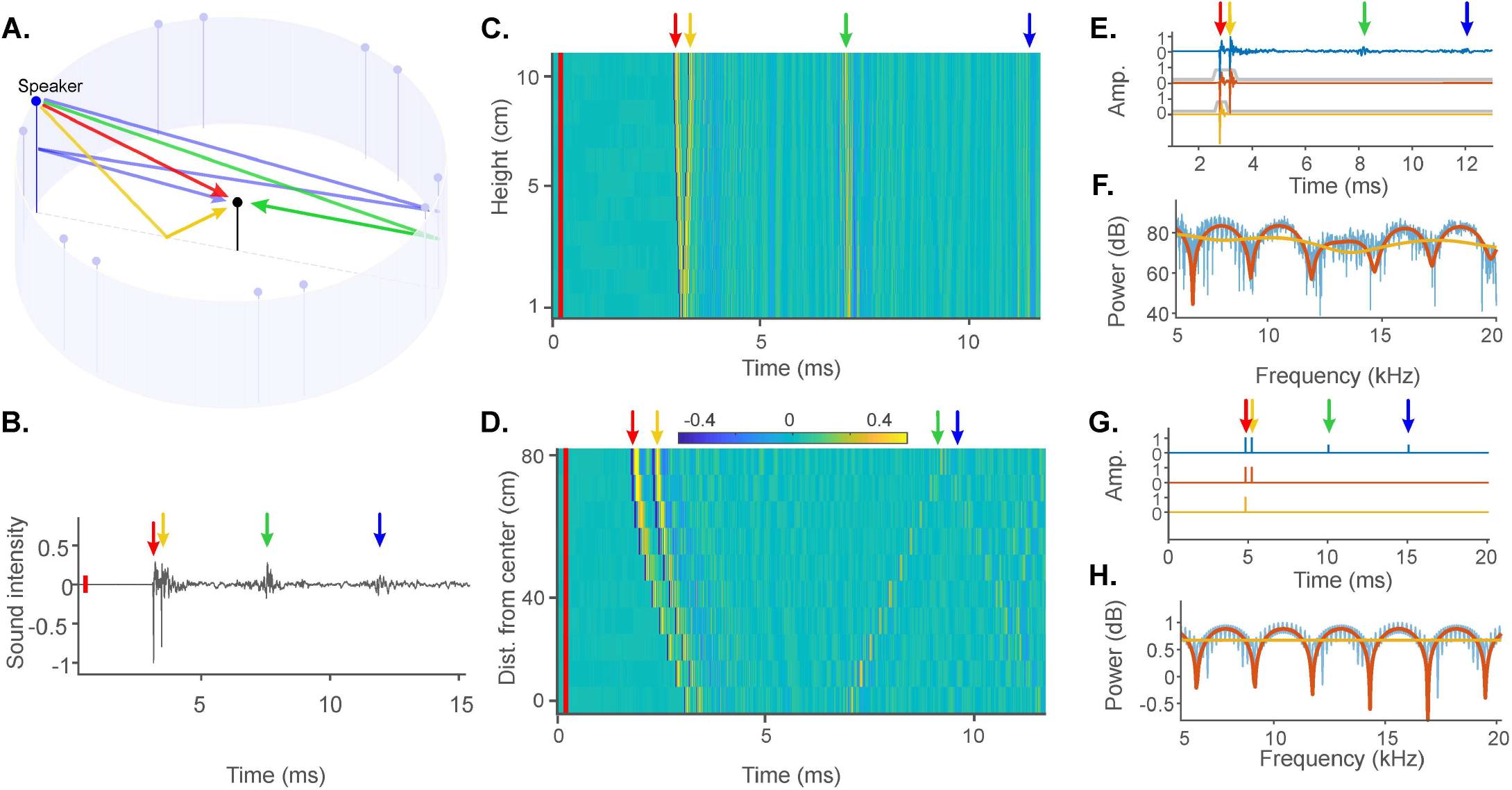
Impulse responses are largely shaped by echoes from the floor and walls. **A**. Schematic view of the echoes inside the arena. Colored arrows represent possible propagation paths of the sound emitted by the speaker and recorded by the microphone (black stem in the middle). **B**. Discrete events, marked by colored arrows, are identifiable in the IR. Red bar at the beginning marks the moment when the impulse would be produced by the speaker. **C**. Impulse responses as a function of height above the floor. Each row represents one IR, colored by amplitude. The microphone was raised 10 times in steps of 1 cm, starting from a height of 1 cm. The colored arrows correspond to the discrete events from B. **D**. Impulse responses as a function of distance along the radius from the center of the arena (bottom) to the wall, just underneath the speaker (top). All IRs were measured at constant height of 10 cm. **E**. An example IR (blue line) was windowed to remove echoes (windows displayed as grey trapezoids). The red version includes only the floor echo, and the yellow version includes only the direct sound. **F**. TFs of the IRs in C. **G**. Approximating the IR as a sequence of four impulses, corresponding to the direct sound and the three echoes. Same conventions as in E. **H**. TFs of IRs from G.

For the IR shown in Fig. 3b, the path length of the direct sound was 0.95 m (red arrow), and the first reflection had a path length of 1.07 m (yellow arrow). This is the expected path length of a reflection from the floor. To illustrate this claim, IRs from one speaker location (speaker no. 5) were measured as a function of the height of the microphone, above a fixed floor location. The difference between the path lengths of the direct sound and the floor reflection increases with microphone elevation (see the geometrical analysis in Supp. Fig. 8). Figure 3c shows 10 IRs measured at a fixed floor location as a function of elevation, stacked from closest to the floor (lowest IR) and to the highest (upper IR). Indeed, increasing microphone’s height decreased the ToA of the direct sound (red arrow, because it decreased the distance between the loudspeaker and the microphone), but increased the delay to the first reflection (yellow arrow). We conclude that the first echo is most likely a reflection from the floor.

The origin of the later two reflections (Fig. 3b, green and blue arrows) was determined using the same approach. Their path lengths in Fig. 3b were 2.39 and 3.98 m, corresponding to 1.5 and 2.5 times the diameter of the arena. These distances are compatible with one or two reflections from the walls (see Fig. 3a for the geometry). In order to verify our hypothesis, we recorded IRs in different locations along the radius of the arena, starting at the center of the arena and moving towards the speaker (Fig. 3d). Both first and late echoes are easily observed in this experiment, and marked on fig. 3d by the yellow, green and blue arrows. The second echo (corresponding to a single wall reflection, green arrow) shows longer delays for microphone positions close to the speaker (when the distance from the far, reflecting wall is the longest) and shorter delays in microphone positions close to the center of the arena. The third echo shows the reverse dependence (blue arrow), also as expected.

Since the IRs are dominated by discrete echoes whose geometrical origin could be clearly determined, we investigated the effect of such discrete echoes on the TF. Generally, a discrete echo causes oscillations in the amplitude of the TF. The relationship between echo delay and rate of fluctuations of the TF can be analytically demonstrated for model TFs composed of only two pure-delay echoes. In that case, a simple calculation shows that the amplitude fluctuations of the TF occur at a rate of 1/Δt, where Δt is the echo delay. These effects are illustrated in Fig. 3e-f. Figure 3e (blue line) replots the IRs in Fig. 3b. We approximated this IR by a sequence of four delta functions, positioned at the onsets of the direct sound and of the three echoes we considered (the floor and the two wall reflections, Fig. 3g, blue). The Fourier transform exhibits a similar profile to the measured one (Compare the blue plots in Fig. 3f, h), with both fast and slow oscillations.

The TF shows slow oscillations with a period of about 2.5 kHz, and rapid ones with a period of about 150 Hz. We isolated the effect of the delay to the first echo on the periodic structure of the TF by windowing the IR to remove the later echoes and observing the effect on its TF. The red plot in Fig. 3e depicts the original IR after all echoes were removed except the first one (originated by the floor), while the red plot in Fig. 3g depicts the corresponding approximation by two impulses. The corresponding TFs (Figs. 3f and 3h, red lines) preserved only slow oscillations. The yellow plots in Figs. 3e-h depict the IR after all echoes were windowed, resulting in a TF that had neither the slow oscillations nor the fast ones.

Figure 4 quantifies these relationships in a different way. TFs recorded at higher positions above ground had faster oscillations than those recorded close to the ground (Fig. 4). This periodicity can be quantified by computing the cepstrum – the Fourier transform of the log amplitude of the TF. This analysis was performed on a frequency range of 3–40 kHz (Fig. 4a), within the frequency range of the speakers. Figure 4 b,d,f displays the TFs recorded at heights 1, 6, 11 cm, respectively, with the corresponding cepstra displayed in Figs. 4c,e,g. These plots clearly show the correlation between microphone’s height above the floor and the periodicity of the TF. Indeed, the time of the peak cepstrum corresponded well with the delay between the direct sound and the floor reflection (0.05 and 0.4 ms, as in Supp. Fig. 8 and Fig. 3c).

**Fig 4.**
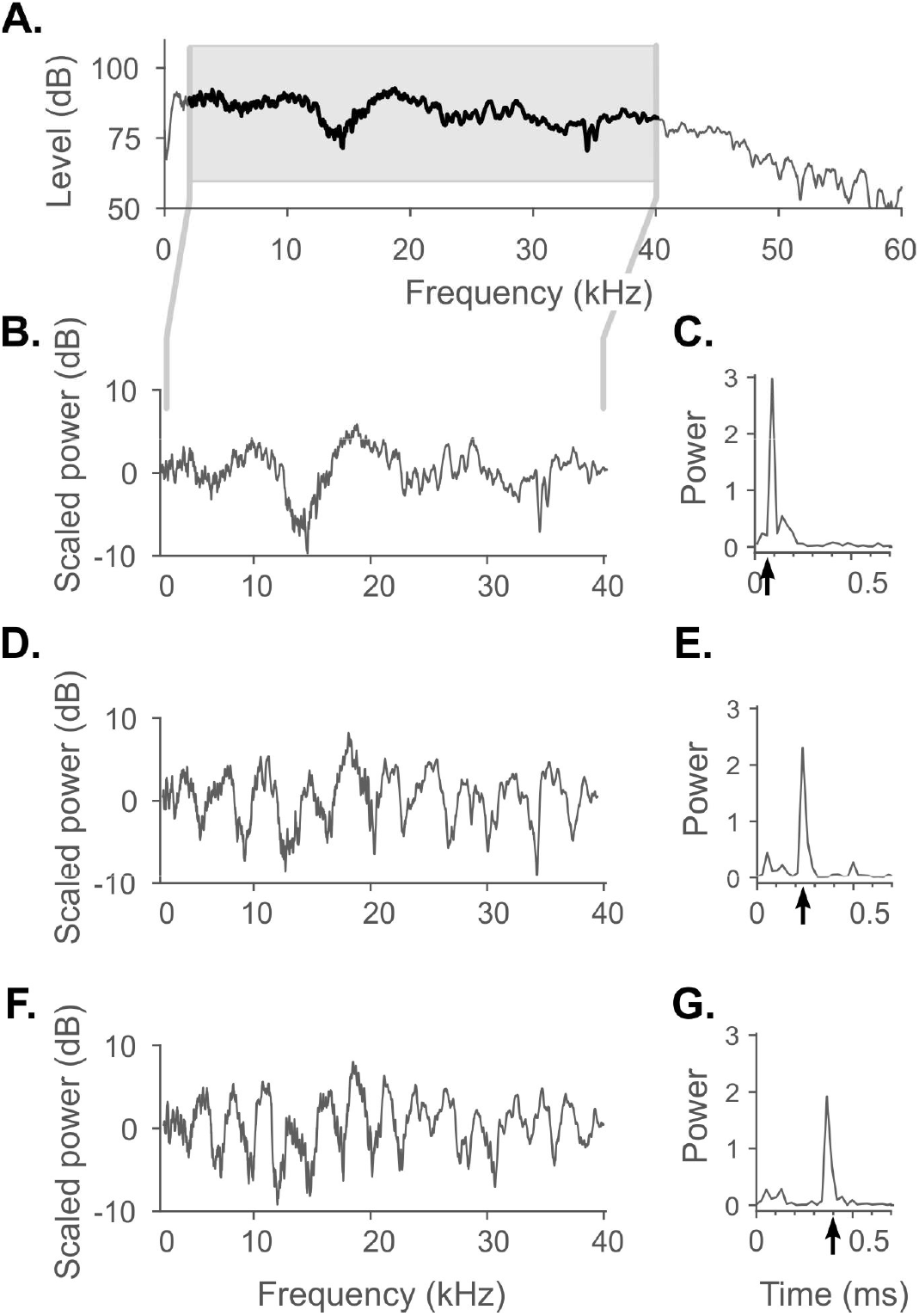
Quefrency (frequency of the oscillations in the TF) increases with height above the floor. **A**. TF was calculated at microphone elevation of 1 cm above the floor. **B**. A frequency region of 0–40 kHz was cropped out and high-pass filtered. **C**. The power cepstrum of B., calculated as |*IFFT*(*log*(*FFT*|(*S*)|^2^))|^2^. The black arrow indicates the measured time delay between the direct sound and the first echo **D**. and **E**. Same as B and C, for a height of 6 cm. **F**. and **G**. Same as B and C, for a height of 11 cm.

We conclude that the first echo in our arena introduces the slow oscillations while the later echoes are responsible for the fast ones. Although the slow oscillations were observable along most of the relevant frequency range (0–60 kHz), their amplitude didn’t exceed 10 dB. The fast oscillations, on the other hand, had much larger magnitude, sometimes spanning almost 60 dB within a few 10s of Hz.

### 3.3 Effect of speaker directivity on the TFs

Speakers radiate sounds non-uniformly in space^13^. Although the TF on the main axis of the speaker may have roughly constant amplitude (as in Fig. 3b), off-axis low frequencies tend to have higher amplitudes than high frequencies. A freely moving rodent is always located off the main axis of at least 10 speakers, making the directivity a factor that is to be considered in the acoustic calibration process.

More formally, typical directivity patterns depend on the size of the speaker through the combination *k* ⋅ *a*, where *k* = 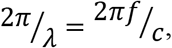, with *f* the frequency, *c* the speed of sound and *a* the radius of the speaker. For *k* ⋅ *a* ≪ 1, the speaker is omnidirectional. For *k* ⋅ *a* > 1, the radiation pattern has a typical shape of 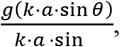 for a *g* which is overall decreasing as a function of frequency, but may show a significant oscillatory behavior as well. Near the axis of the speaker, θ is small and the dependence on frequency is therefore weak. Farther away from the axis, the directivity pattern causes a frequency dependence which becomes stronger as the angle θ increases, and whose details depend on the exact function *g*.

These directivity properties are thoroughly explained by Müller & Möser (2013)^14^ and Geddes (2009)^15^. To evaluate the directivity of our speakers (since directivity specifications were not supplied by the manufacturer) we measured TFs for different angles θ relative to speaker’s main axis (Fig. 5). The TFs were measured at 23 locations along a 50 cm circle around the speaker location. These 23 locations spanned a central angle of 120º around the speaker, one point on the principal axis and 11 on each side.

We subtracted the TF at the point that coincides with the principal axis of the speaker from all other TFs to create relative TF profiles, shown in Fig. 5a. This measure preserves the relative attenuations along the different angles while removing location-independent distortions created by the imperfect audio equipment (loudspeaker and microphone). We compared these measurements with the expected directivity pattern of a disk vibrating in free space (Eq. 2, Fig. 5b), where the function g is known to be *J*_1,_ the Bessel function of the first kind (up to a numerical factor). *c* was the speed of sound at 20 C°, 343 m/s, and *a* the effective speaker radius (1.8 cm):

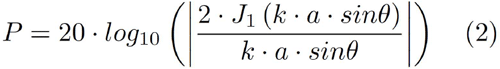

These plots display the sound power as a function of both the frequency and the angle from the speaker’s main axis. For the range of angles that we measured, frequencies below 15 kHz had an almost constant power. This is further visualized in Fig. 5c, where all the positions of the microphone are displayed, colored by the sound power. At higher frequencies, however, the sound is more attenuated off-axis than on-axis (Figs. 5d,e). We conclude that to avoid acoustic variability at high frequencies, the speaker should be selected so that the rat is as close as possible to its acoustic axis.

### 3.4 Narrow-band noise stimuli reduce the effects of fast fluctuations in TF amplitude

The most problematic feature of the TFs identified above consists of the fast fluctuations in amplitude, which may be very large (dozens of dBs). We identified them as the consequence of the wall reflections, of which the earliest would occur at a delay on the order of the travel time along a diameter of the arena, which is about 5 ms. Thus, the slowest of the fast fluctuations occur over bandwidths of 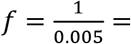 200 Hz. Longer delays would result in faster amplitude fluctuations.

We show here how to reduce the effects of the fast fluctuations on sound amplitude through the use of narrow band stimuli instead of pure tones. Pure tones are widely used in auditory studies due to their simplicity, but in echoic environment their amplitude varies substantially from place to place as well as for different frequencies because of the fast fluctuations in the TFs. We generated narrow band stimuli as combinations of a large number of closely-spaced pure tones, superimposed at random phase. We reasoned that this superposition would average out the rapid fluctuations in the TFs, making sound level much less position- and frequency-dependent: at any location in the arena, some of the pure tone components would be strongly attenuated and other won’t, so that the overall power of the narrow band stimulus should be less variable than that of the individual pure tones composing it. The extent of this averaging can be controlled by the frequency span of the narrow band noise, which we set to 0.5 kHz (2.5 times the width of the typical fluctuation).

**Fig 5.**
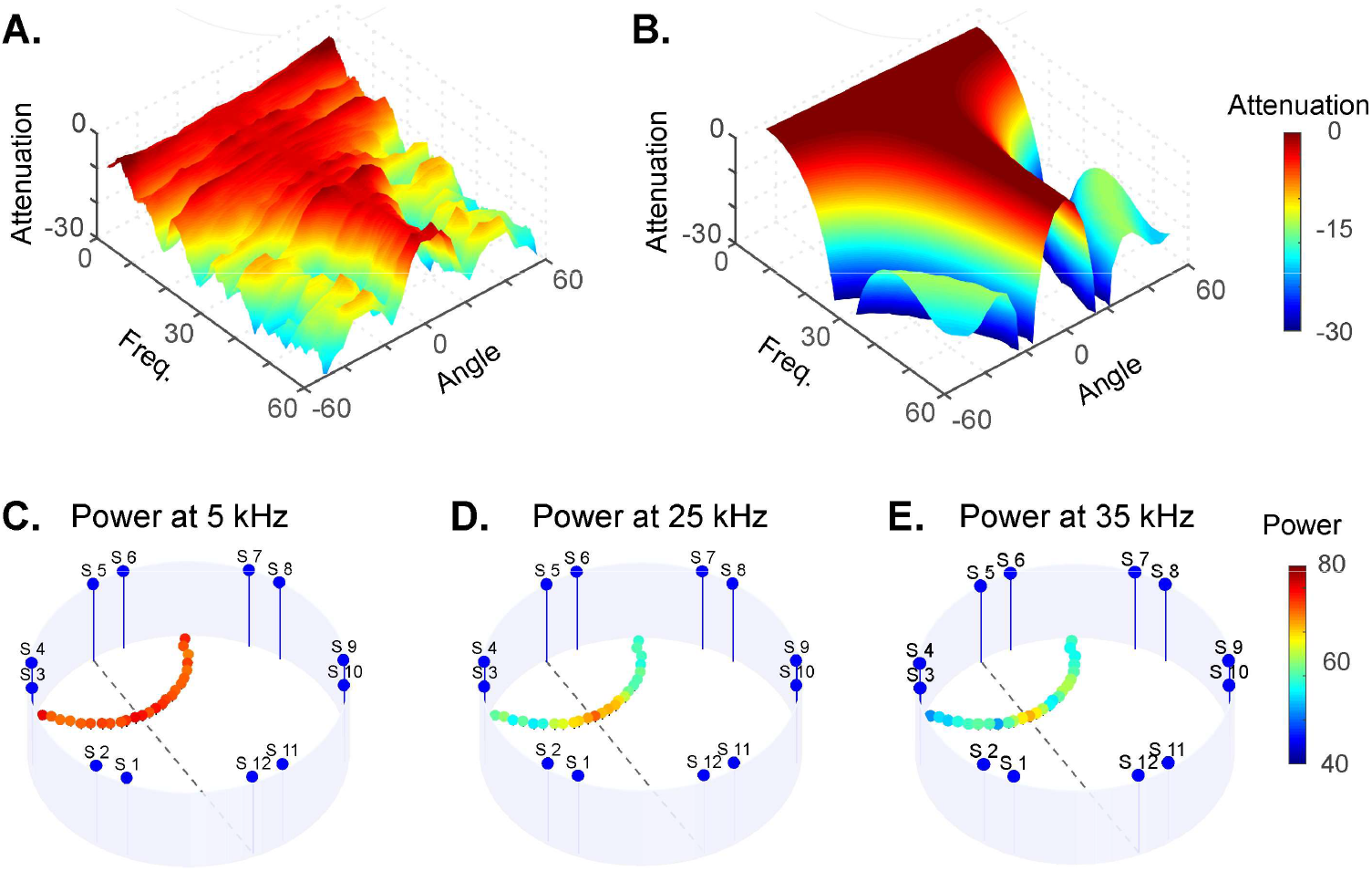
Speaker directivity. **A**. TFs were sampled at 23 locations, 11 at each side of the main speaker’s axis. All samples maintained a constant distance to the speaker (No.5). All TFs were stacked together and colored the sound power, then the TF at the central point was subtracted from each of them to produce the relative TF profile. **B**. Expected directivity for an oscillating round piston with the same radius as the speaker. **C**. Relative power at 5 kHz at all sampled locations. **D**. Same as C., at 25 kHz. **E**. Same as C, for 35 kHz.

To assess the efficiency of the proposed method, we first recorded responses to a 10 s long frequency sweep spanning the range of 1 – 1.5 kHz, and verified that the amplitude fluctuations of the recorded signal (over a range of about 20 dB) were essentially equivalent to the power fluctuations of the TF measured using the Golay codes (Supp. Fig. 9). We then compared the envelope fluctuations of a slow pure tone frequency sweep and of a slow harrow band noise frequency sweep (see Methods for the details of the stimuli; Fig. 6). The envelope of the pure tone sweep significantly fluctuated as the frequency changed over a 500 Hz range, due to the fast amplitude fluctuations in the TF. The pattern of the fluctuations varied as a function of the location (Figs. 6a and 6b illustrate recordings at two locations 20 cm apart) and the frequency band (upper plots represent frequency sweep at 0.5–1 kHz, bottom represent 10–10.5 kHz, where the fast fluctuations are less apparent). As expected, narrow band stimuli had substantially lower amplitude fluctuations compared to pure tone sweep.

**Fig 6.**
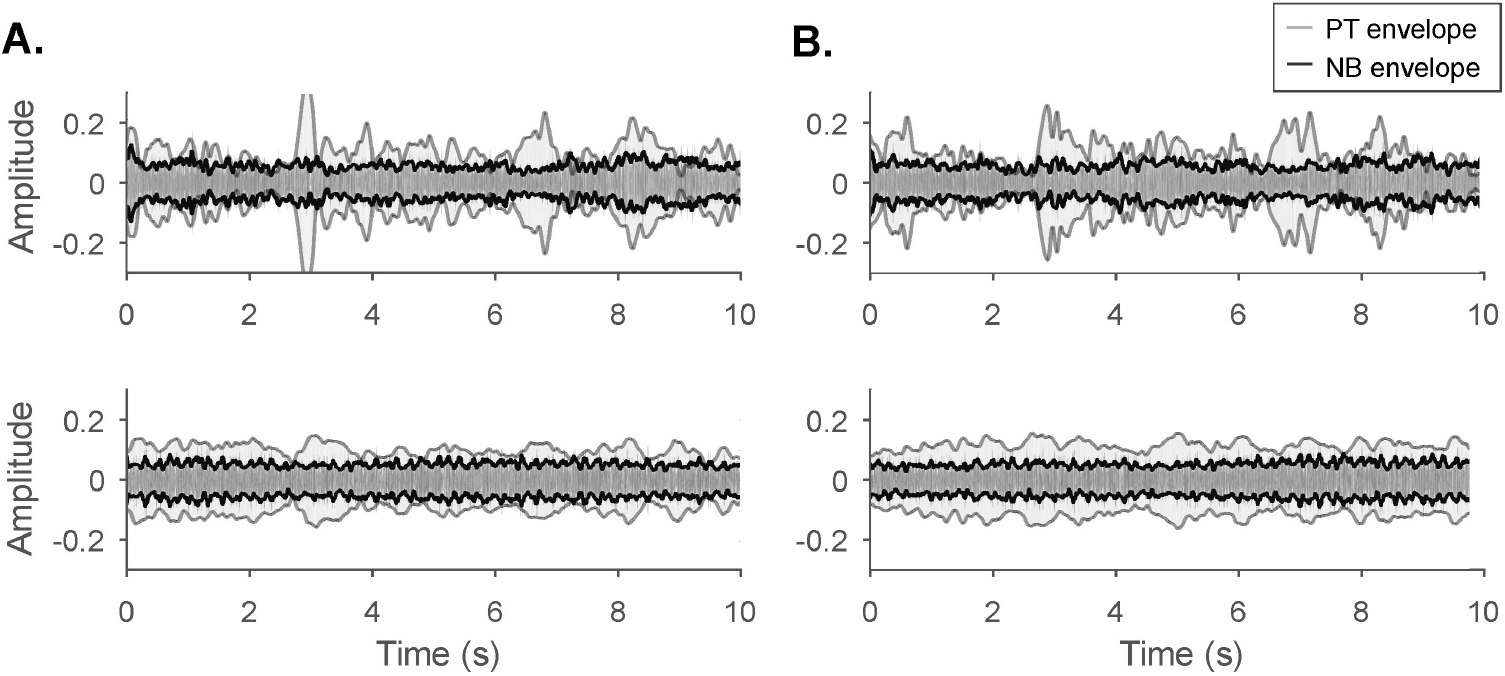
Narrow-band noise has lower amplitude variability than pure tones. **A**. Amplitude fluctuations as a function of time during a slow sweep of a tone (light grey) and a narrow noise band (dark grey and black). Both stimuli covered the same frequency range (1–1.5 and 10–10.5 kHz for the upper and bottom plots, respectively), played by same speaker, and recorded at the same location. **B**. Same as A, at a different location.

Unfortunately, the same approach cannot work for the slow fluctuations in the TFs, caused by the floor reflections. These fluctuations are much slower – the notches vary in their periodicity, from two periods between 0–60 kHz when the microphone is 1 cm above the floor up to dozens of periods when the microphone is 10 cm above the floor (Fig. 4). One approach for minimizing the amplitude modulations across the arena, consists of using an average TF from many different locations and speakers as a guide for a robust stimulus generation. To demonstrate this approach, we averaged 4680 TFs that were sampled from all 12 loudspeakers at 390 locations (Supp. Fig. 10a), and looked for a wide range of frequencies that had minimal power span. Supp. Fig. 10b depicts this process and shows the resulting five narrow band stimuli picked at 7.5 kHz steps along a range of 30 kHz. The recorded amplitudes of these stimuli spanned only 10 dB range.

### 3.5 Acoustic localization of freely moving animal in 3D

The traveling distance of the direct sound can be evaluated with precision of up to 0.3 cm (Results section B.). Given 3 distances from loudspeakers to the microphone, the later can be localized by numerical solution of the multi-lateration problem (see Methods and Supp. Fig. 6). If the microphone is carried by an animal, it can be used for 3D tracking as the animal freely moves through the environment. We tested this approach with freely moving rats that were carrying a headstage containing a microphone.

Implementing this method requires solving two problems: First, the acoustic quality of the headstage microphone (denoted by HS) was inferior to the calibrated B&K microphone (BK) we used for measuring the TFs. Second, the rat may move while the calibration sounds are presented, introducing inconsistencies in the distances used for multi-lateration. Both these factors reduce the localization accuracy. We therefore tested the accuracy of the method, varying the duration of the Golay pair (and therefore the SNR of the estimated IR) as well as the number of loudspeakers used for estimating the animal location. We performed these measurements on a static microphone as well as on a freely-moving rat carrying a headstage containing a microphone, verifying the localization of the rat using a concurrently collected video movie.

The precision of the trilateration solution was assessed for statically placed microphones. First the acoustic localization was compared to manually determined location of the microphone, showing that the acoustic 3D localization is consistent with the true location of the microphone (Fig. 7a-b). Then ten repeats of the localization process were performed, each based on distances to all 12 loudspeakers (Supp. Fig. 11a, blue dots). The localization error was evaluated as the distance of the solution point from the cluster center, indicating the cluster volumetric spread. The mean errors were 0.061 cm and 0.099 cm for the BK and HS microphones respectively.

**Fig 7.**
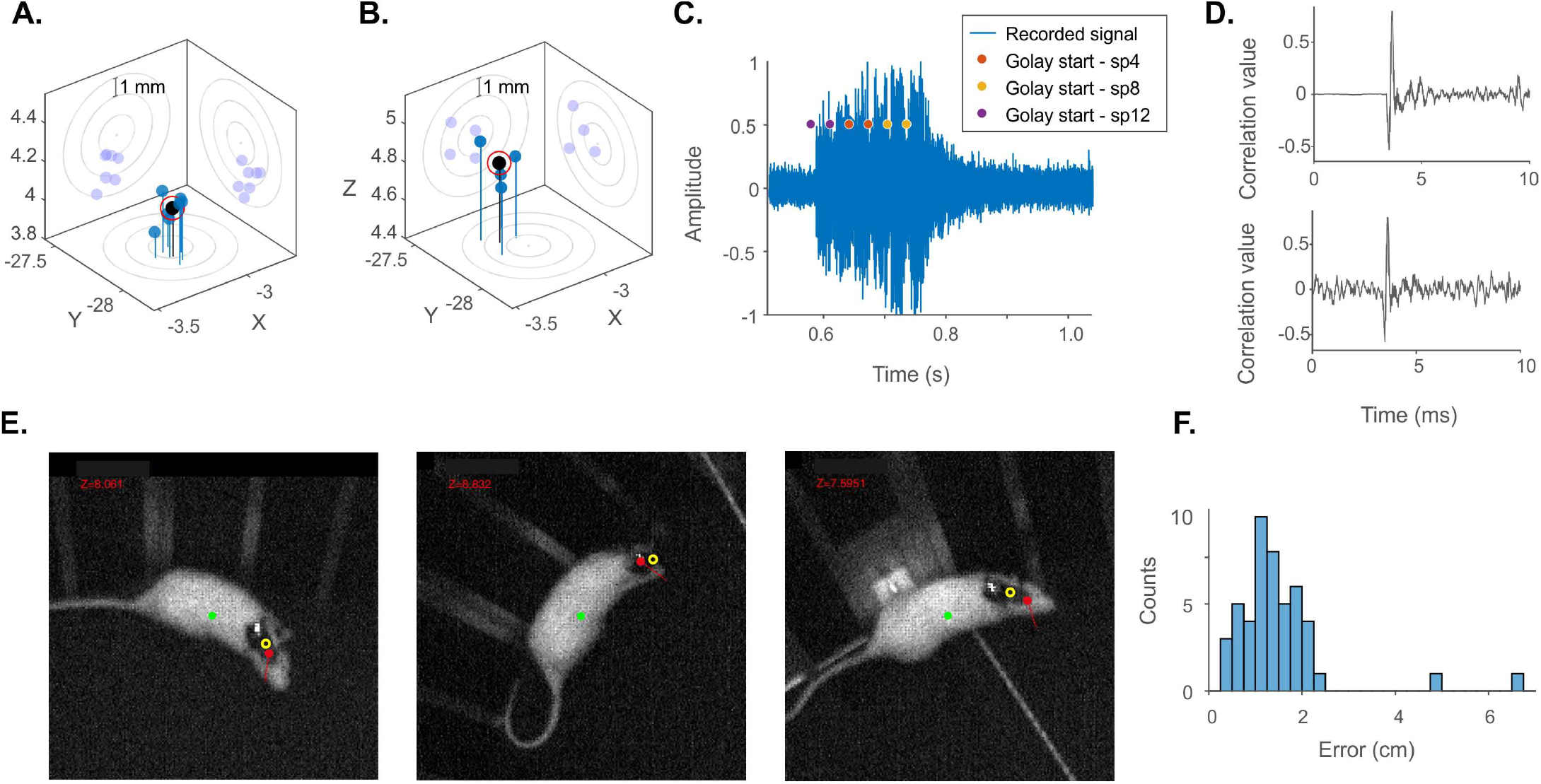
Acoustic 3D localization of a freely moving animal. **A**. A statically placed microphone was acoustically localized using the delays to the direct sounds emitted by 12 speakers (Golay codes of length 2^16^ bits, 10 independent localizations marked by blue stems). The location of the microphone was manually evaluated with uncertainty error of 0.3 cm (projected to the three 2D planes as concentric circles). The mean spread of the localizations was 0.085 cm from the cluster center (black stem with red circle). **B**. Same as A. but for acoustic localization using 3 speakers, resulting in mean error of 0.12 cm. **C**. A pair of Golay codes (10^12^ bits) are played by 3 different speakers (marked by colored dots) with 10 ms silence buffer in between. This sequence is recorded by the microphone as a continuous sound (blue plot) and is used to locate it in 3D. **D**. The precision of the localization depends on the sound to noise ratio of the IR, that is controlled by length of the Golay and the silence buffer. When both are long enough, the SNR is high (upper plot). Bottom plot displays an example of an IR with low SNR. **E**. Frames from Supp. Movie No.2 (Golay of length 10^12^ bits, silence buffer 10 ms). The green dot is the centroid of the rat as registered by the camera, the yellow circle is the manual annotation of the location of the microphone and the red stem is the location of the microphone estimated acoustically, projected to the 2D image. **F**. Mean localization error of Supp. Movie No.2 is 1.5 cm (based on 50 measurements). The error is defined as a distance between the acoustic localization of the microphone, projected to the 2D image (the red point in E.) and the manual annotation (yellow circle).

The superior precision of BK can be explained by the higher sampling rate we used for these measurements. Sampling at 192 kHz provided a resolution of 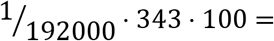0.18 cm, while the sampling rate of the HS was limited by hardware constraints to 44 kHz, resulting in a resolution of 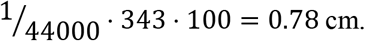 Another reason could be the inferior recording quality of HS, as shown in the strongly filtered TFs that were recorded by HS (TFs recorded at the same location of the arena are compared in Supp. Fig 12b). In consequence, the IRs estimated from HS had slower rise of the IR, as illustrated in Supp. Fig. 12a around 0.7 ms, complicating the detection of the onset latency and reducing localization accuracy. For both BK and HS, the 3D localization error was a few times smaller than the distance resolution. We believe this decrease is the result of combining the twelve distances during the multi-lateration process.

We studied the precision of the localization as a function of the parameters of the sound presentation protocol. A localization protocol consisted of selecting the speakers from which the sounds would be presented, as well as the durations of the Golay code and of the silence buffer. For example, Fig. 7c shows the sound recorded during a localization protocol consisting of 3 speakers, each playing a pair of Golay codes. Each Golay code was 2^12^ bits long (sampled at 192 kHz, thus lasting 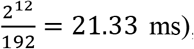, with a silence buffer of 10 ms in between. As already mentioned, the precision of the determination of the speaker-microphone distance depends on the precision of the IR onset. The precision of the IR onset mainly depends on two factors: The power of the calibration sound (regulated by the Golay code length, in bits) and the duration of the silence buffer between successive Golay sequences (within and across pairs): this duration determines the perturbation caused by one Golay sequence on the next one. Thus, there is a tradeoff between the SNR of an IR and the length of the localization cue. For example, using Golay sequences of length 2^16^ bits and silence buffer of 0.2 sec resulted in clear IR onset (Fig. 7d, top) while using Golay sequences of length 2^10^ bits and silence buffer of 0.01 sec resulted in substantial fluctuations around the IR onset, masking the exact onset (Fig. 7d, bottom; Supp. Fig. 13).

The duration of the localization process can be furthermore shortened by reducing the number of speakers. While three speakers are enough to define a trilateration problem, adding more speakers would improve the precision. However, each additional speaker prolongs the localization process. For example, trilateration based on 3 speakers requires only 3 pairs of Golay codes to be played while trilateration based on 12 speakers is more than 4 times longer (Supp. Fig. 14). We tested the effect of the speakers set size on microphones at a fixed position, by using only 3 speakers for the localization of a static microphone (Supp. Fig. 11b). The localization error grew to 0.21 cm and 0.44 cm for BK and HS as expected, but maintained sub-centimeter precision. We concluded that three speakers are enough for precise localization.

Lastly, we tested the acoustic localization on a freely moving rat. We tried four localization protocols that differed in Golay and silence buffer lengths, as well as the number of speakers (listed in Supp. Fig. 15 with the corresponding localization precision). Each protocol was comprised of 50 localization trials, played once every 2 s, and the signal recorded by HS was stored for offline analysis. Additionally, a visual localization was performed using a camera. The acoustic localization error was determined as the distance from the acoustically reconstructed location of the microphone (Red dot in Fig. 7e., Supp. movies 1–4) to the manually annotated location of the headstage (empty yellow circle; see Methods). The highest accuracy was obtained for a protocol using 3 speakers, a Golay code length of 2^12^ bits, and silence buffers of 0.01 s, resulting in a mean error of 1.5 cm even for fast movement bursts (Fig. 7e,f and Supp. Movie 2). Shorter Golay codes (of 2^11^ bits) led to increased mean error, reaching 4 cm on average (Supp Fig. 15a.), significantly worse than similar localization cue with Golay of length 2^12^ (Supp. Fig. 15b). Similarly, using more than 3 speakers led to decreased precision (Supp. Fig. 15c, d), presumably due to prolonged localization protocols.

## 4 DISCUSSION

We used Golay complementary sequences to estimate impulse response (IR) of the arena^5,6^. We demonstrate a principled analysis of the IRs, identifying the geometric basis for their structure, the resulting distortions in the frequency domain, and suggesting approaches for minimizing the acoustic inhomogeneities as a function of frequency and spatial location. We then demonstrate a viable, high precision localization of the animal using the same hardware used to calibrate the arena.

### 4.1 Automated calibration pipeline

This study introduces a pipeline for acoustic calibration of echoic environments, which was successfully applied to our own setup. The initial step of the sound sampling is a laborious task, if performed manually: Sound samples are to be collected at high spatial resolution along few meter-wide environment, due to high variability of perturbations between two adjacent locations. Moreover, the IR has to be calculated individually for each speaker, increasing the recordings count to hundreds. Clearly, a fully automated sampling system is the only way to address this challenge. We achieved it using a programmable robot that performed the sound sampling routine, autonomously operating for hours over hundreds of predefined recording locations.

### 4.2 Characterizing the acoustic properties of the arena

The IRs had relatively simple structure, consisting of a direct sound, a short-delay floor reflection, and longer delay wall reflections (Figs. 2, 3). We showed that both floor and the wall echoes were clearly visible on the IR, across most locations and speakers, and that these reflections determined much of the spectral distortions in the TFs. Importantly, we did not detect echoes from sources outside the arena, including the ceiling. The ceiling is located 2.1 meter above the arena’s floor, reflecting echoes with approximate traveling path of 4.2 m, corresponding to ToA of 12–13 ms. Perturbations are indeed visible on the IR at about 12 ms (Fig. 3b, blue arrow), but we identified them with the second wall reflection, which also had ToA of 12 ms. The absence of a ceiling reflection could be due to the directivity of the loudspeakers: the ceiling was approximately located at 125 degrees off the acoustic axis, and the amplitude of the sound could be substantially lower at the direction even at the lowest frequencies of interest. This observation supports the simplifying assumption that the acoustic effects of external objects are negligible, and the arena can be modeled as an environment with only two sources of echo – the floor and the wall.

Approximating the IR as a sequence of echoes made it easy to understand the structure of the spectral distortions: echoes cause periodic fluctuations in the amplitude of the TFs, with the time intervals between echoes determining the rate of these fluctuations. Short time intervals resulted in slow fluctuations, while long time intervals resulted in rapid fluctuations in the TF (Fig. 3e-h).

We believe that one reason for the relatively simple structure of the IRs is the use of a large, circular arena. This caused a clear separation in time between the floor and wall reflections. Since we showed that the effects of the wall reflections can be largely neutralized through the use of narrowband stimuli, further optimization of the arena for acoustic purposes would consist of reducing the floor reflections. This can be achieved by using absorbing or dispersing materials on the floor, although this would complicate the maintenance of the arena.

An effective way of minimizing the distortions is a guided design of the auditory stimuli. It is advisable to use mostly frequencies below about 15 kHz to avoid the influence of speaker directivity (Fig. 5), to use narrow band stimuli instead of pure tones (Fig. 6) to reduce the effects of wall reflections, and to be guided by an average TF profile as in Supp. Fig. 10.

The calibration pipeline may be extended for generating narrowband stimuli, based on the following steps: Record IRs across the arena and compute their TFs, calculate the frequency of the fast spectral notches (the quefrency peak, as in fig. 4), generate the narrow band stimuli and measure their amplitude as a function of center frequency in order to evaluate their performance. These stimuli can provide best fitting acoustic cues for the unique geometry of the specific setup.

### 4.3 Application of the acoustic localization of freely moving animals

We developed a 3D acoustic localization technique that provides a spatial resolution of about 1.5 centimeter in 160 cm wide environment. Hardware pre-requisites of the acoustic localization are three speakers, a microphone, and synchronized clocks of the sound reproducing and recording equipment. It is worth noting that the presented method particularly fits experimental setups that possess integrated microphones on the head stages or body harnesses of the animals.

There are a number of advantages of the method over alternatives: Its hardware pre-requisites are relatively modest, especially beneficial for those setups that already utilize acoustic equipment; The localization is performed in a fraction of a second in a closed room, due to the small time interval that is required for sounds to cross a few meter environment; It has no footprint on the electrophysiological recordings, since no electro-magnetic disturbance is introduces; and it acquires all three volumetric coordinates (X, Y, Z).

The main disadvantage of 3D acoustic localization is the need to use audible sounds that may interfere with the acoustic paradigm of the scheduled experiment, or may generally stress the animal. It also has a relatively low temporal resolution, limited by the durations of the Golay code and the silence buffer. The temporal resolution achieved here is limited to about 5 Hz.

## 5 ACKNOWLEDGEMENTS

The authors thank Dr. Maciej M. Jankowski and Ana Polterovich for awake rat recordings. This work was supported by a European Research Council (ERC) advanced grant GA-340063 (project RATLAND), by F.I.R.S.T. grant no. 1075/2013 from the Israel Science Foundation to IN, and by a grant from the Gatsby Charitable Foundation.

---

**Supp. fig 1:**
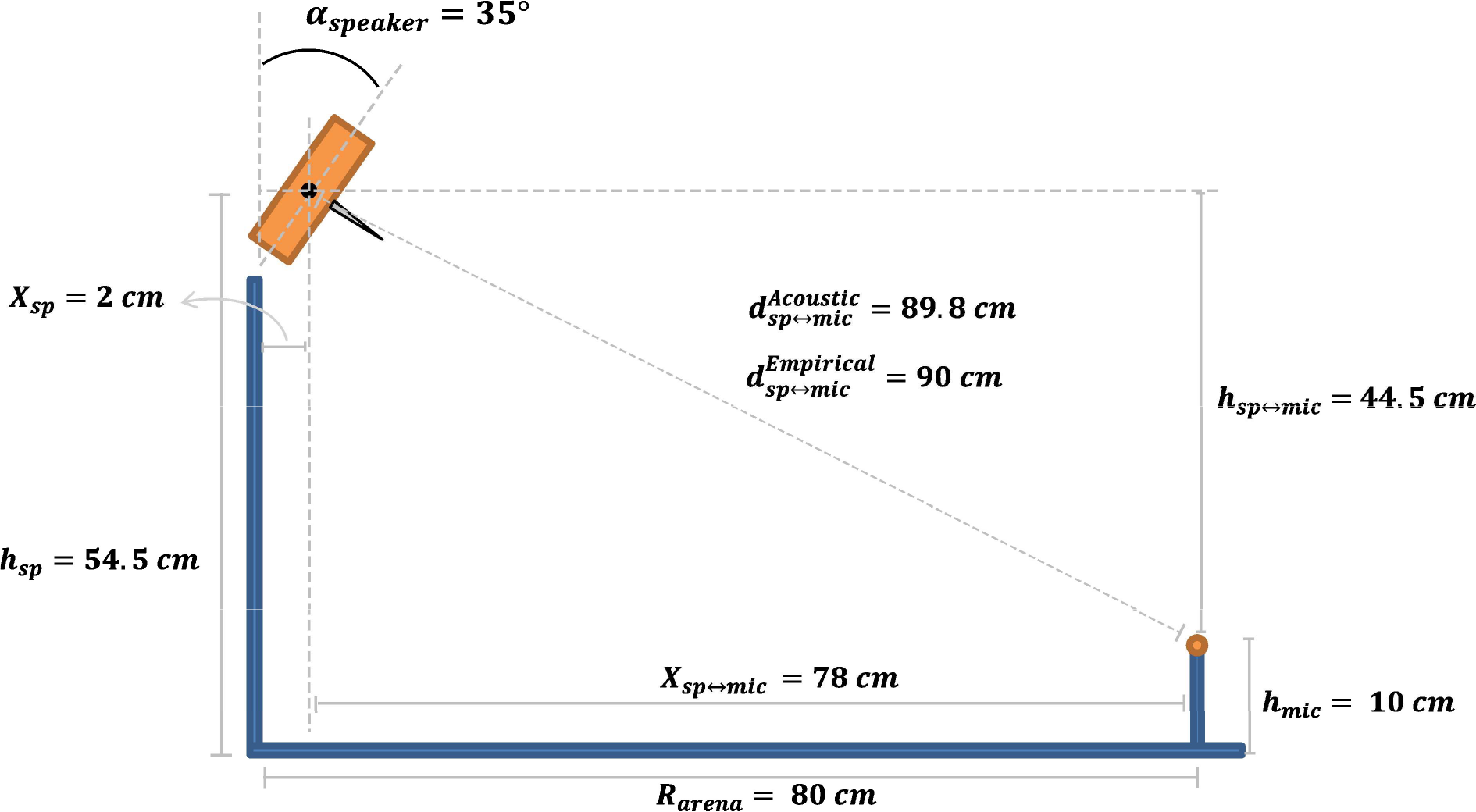
Control for the precision of acoustically evaluated distances. A side view of the environment (speaker – orange rectangle, microphone – orange circle). The manually measured distances: *R_arena_* is the radius of the arena. *X_sp_* is the distance of the speaker’s membrane from the wall, due to the 35° tilt. *X_sp↔mic_* is the horizontal distance of the microphone from the loudspeaker. *h_sp_* is the height of the speaker’s membrane center. *h_mic_* is the height of the microphone. *h_sp↔mic_* is its vertical distance of the microphone from the speaker. Using the Pythagorean theorem we calculate 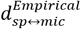, the distance between the speaker’s membrane and the microphone, based on the manual measurements. 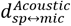 is the acoustically evaluated distance (89.8 cm), deviating 0.2 cm (0.22%) from the manually measured one (90 cm).

**Supp. fig 2:**
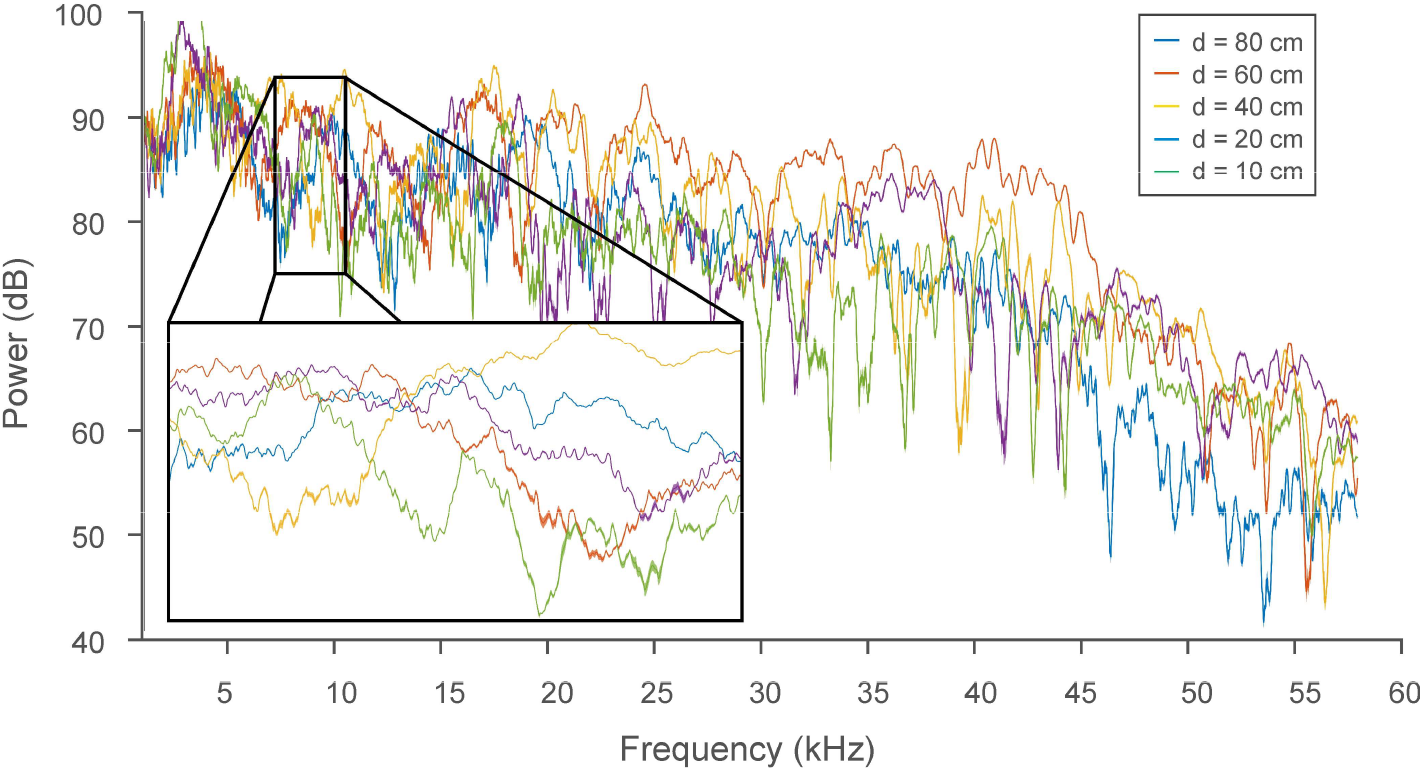
TFs are highly variable between different locations of the arena, but is highly reproducible at each location. TFs were measured in five different locations along a radius connecting the speaker and the center of the arena (d=80 cm). At each location three measurements were made. Each line represents TF, with the shaded area representing the standard deviation of the 3 measurements. The variability between measurements in the same location is hardly visible when compared with the between-location variability.

**Supp. fig 3:**
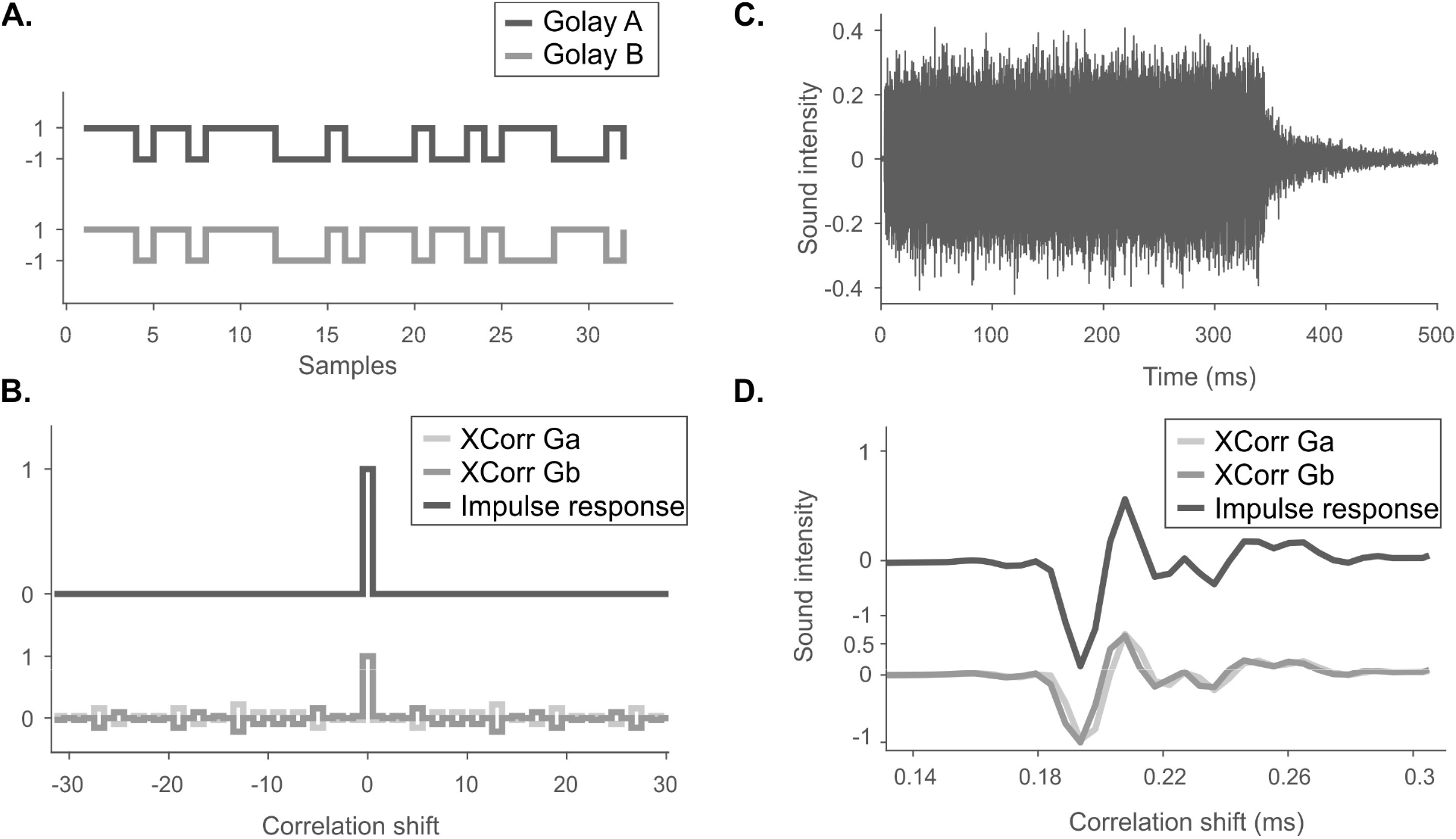
Computing an impulse response from the responses to a pair of Golay complementary sequences. **A**. An example of two complementary Golay sequences of length 2^5^ = 32 *bits*. **B**. The two auto-correlations of Golay codes (bottom two lines) sum to an impulse at the origin. **C**. Sound recording of one Golay sequence. **D**. The responses to each Golay sequence is cross-correlated with its original Golay sequence, producing a single cross-correlation. The two cross correlations are shown by bottom plots. When summed, they form an impulse response (the upper dark line).

**Supp. fig 4:**
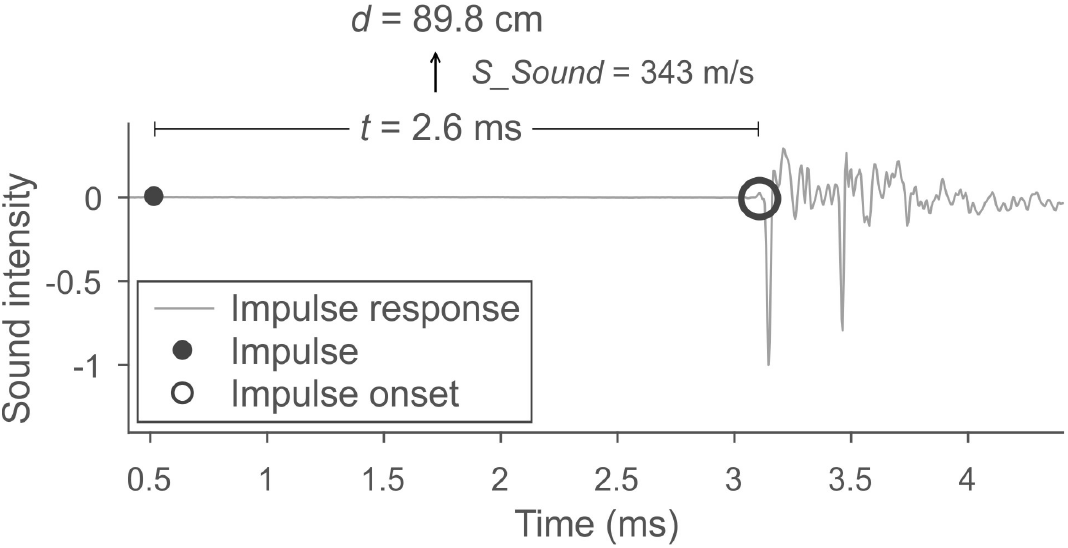
Impulse response encodes the distance of the speaker to the microphone. An impulse response was recorded at the center of the arena (gray line). Black point represents sound onset, black empty circle is the estimated sound onset at the microphone’s location. Time difference of these two points is called time-of-arrival of the sound, in this case it is 2.62 ms. It can be converted to distance by multiplication with the speed of sound: 0.00262 ⋅ 343 = 89.8 *cm*;.

**Supp. fig 5:**
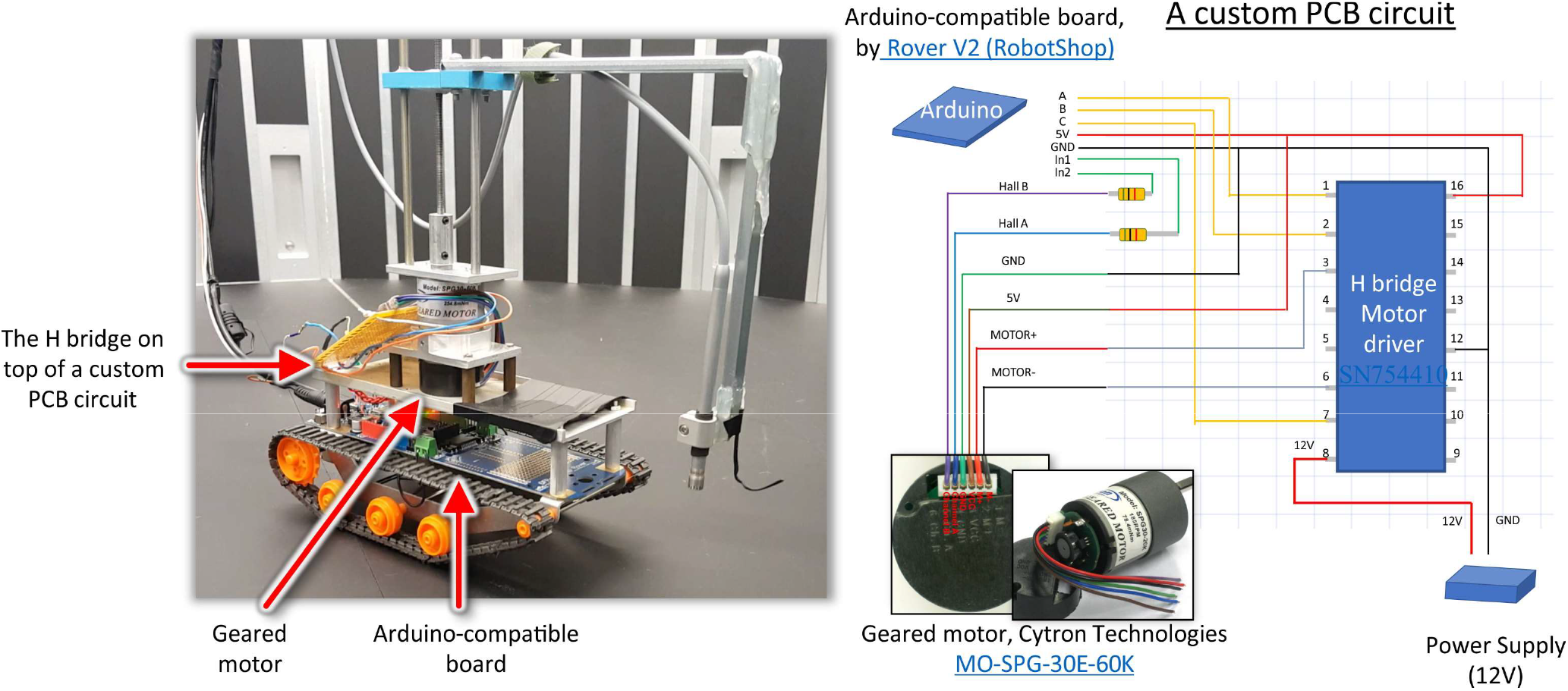
Electrical circuitry that enables the vertical movement of the microphone. The bottom part of the robot was assembled from the kit <u>Rover V2 (RobotShop)</u>. A geared motor (<u>MO-SPG-30E-60K</u>) was installed on top of it, fixed inside a custom made frame. The motor controlled the elevation of the extension arm, that held the microphone. An Arduino-compatible board (part of the RobotShop kit) controlled the robot’s tracks and the geared motor. The later was connected to the board using a custom made PCB circuit, utilizing an H bridge driver.

**Supp. fig 6:**
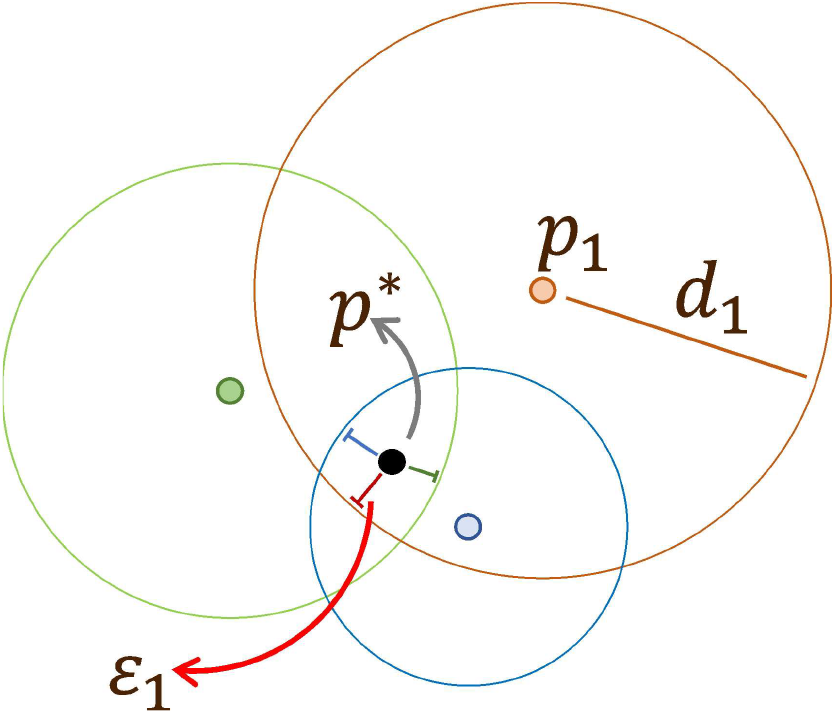
The trilateration for microphone localization in 3D. Trilateration is distance-based localization that relies on intersection of spheres (or circles, for 2D). For acoustic localization, the point of interest is the microphone (black point, p*). The locations of the loudspeakers are known (colored dots, i.e. p_1_) and distances to the microphone are evaluated using the time-or-arrival of the direct sound, estimated from the IRs (i.e. d_1_). The three circles don’t intersect in a single point due to measurement noise, and the location of the microphone is approximated through minimization of distances to the three circles (i.e. ɛ_1_).

**Supp. fig 7:**
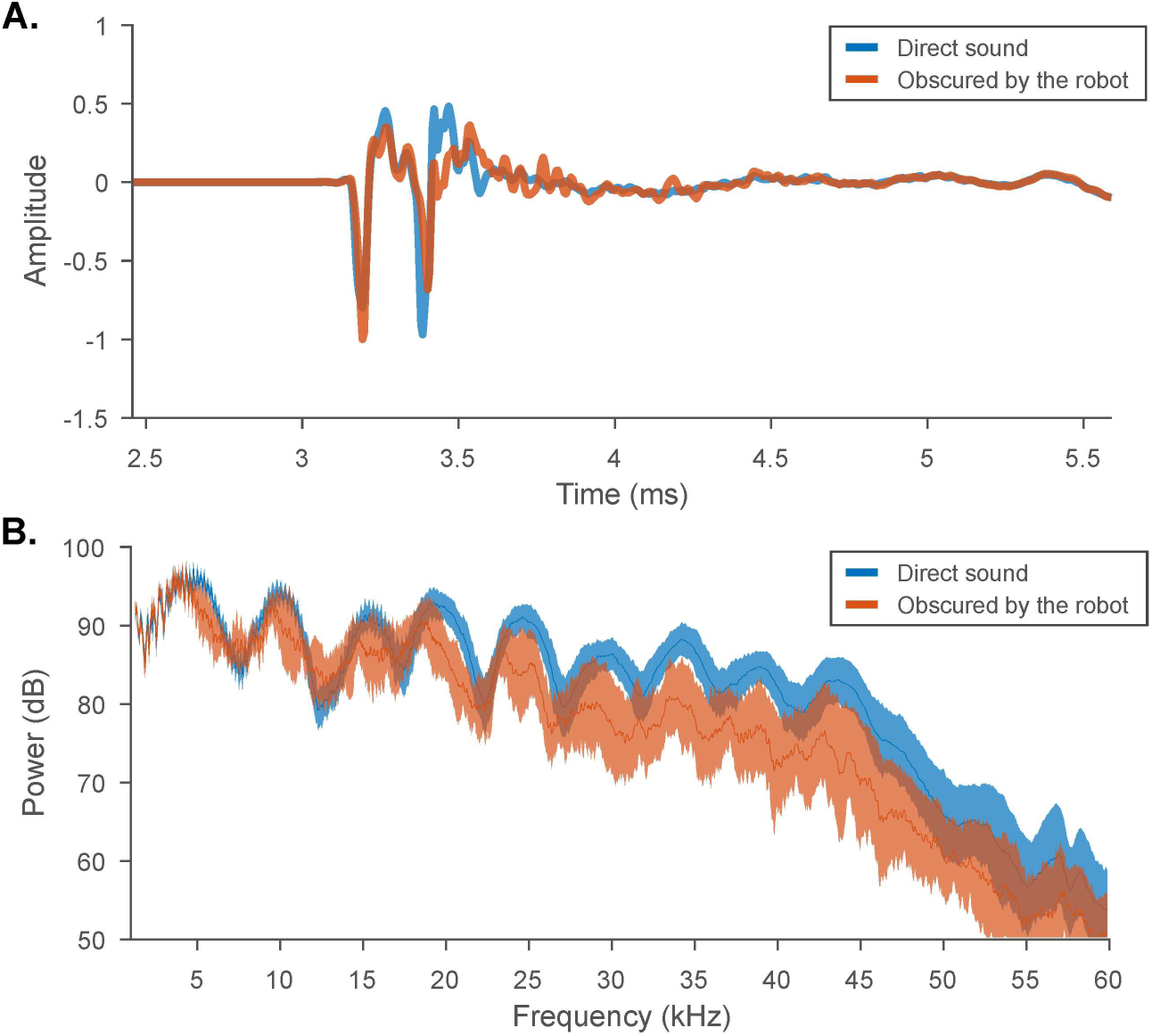
The IR and TF are affected by the robot’s body. The rover was rotated 12 times around the microphone (the location of which was kept constant), each time by 90° clockwise (completing three full circles). After a single rotation, the IR was sampled from all the 12 speakers, out of which two speakers had no line of sight with the microphone because of the robot’s body. **A**. The average IR for all speakers from all body positions that allowed a line of sight with the microphone (blue) and the average IR for the speakers obscured by the robot (orange). Solid line is the mean, shaded areas are the standard deviations across all corresponding robot body locations). **B**. TFs of the impulse responses from A.

**Supp. fig 8:**
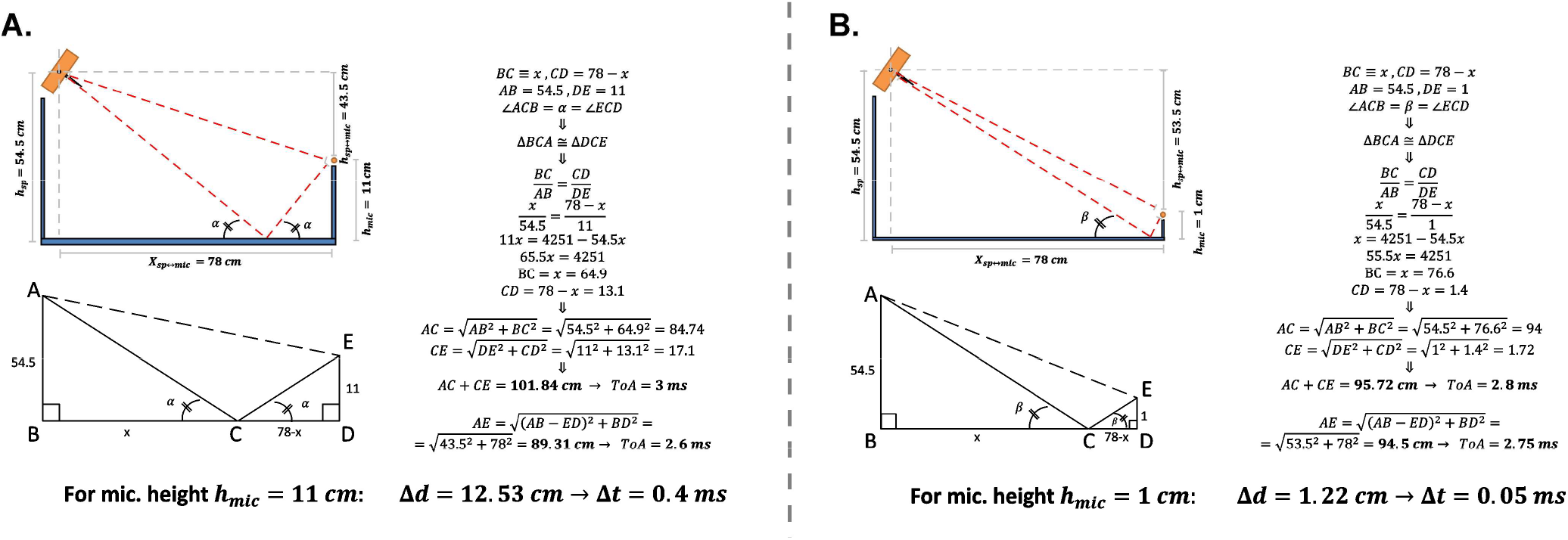
Computing path differences for the direct sound and the floor emitted echo, at the lowest and highest sampled points (*h_mic_*=1 cm, *h_mic_* =11 cm). **A**. The upper diagram shows a side view of the arena. Orange rectangle represents the loudspeaker, orange circle is the microphone that was placed 11 cm above the ground. The floor reflection was computed under the assumption that the incidence and reflection angles are equal. The bottom diagram is a geometric problem arising from the upper diagram. Distances are converted to time through the relation 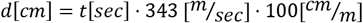 The path difference was found to be 12.53 cm, corresponding to an expected 0.4 ms delay of the echo after the direct sound onset. **B**. Same as A for a microphone placed at a height of 1 cm. Here the path difference decreases to 1.22 cm and the delay to 0.05 ms. These predictions are consistent with the experiment of Fig. 3c.

**Supp. fig 9:**
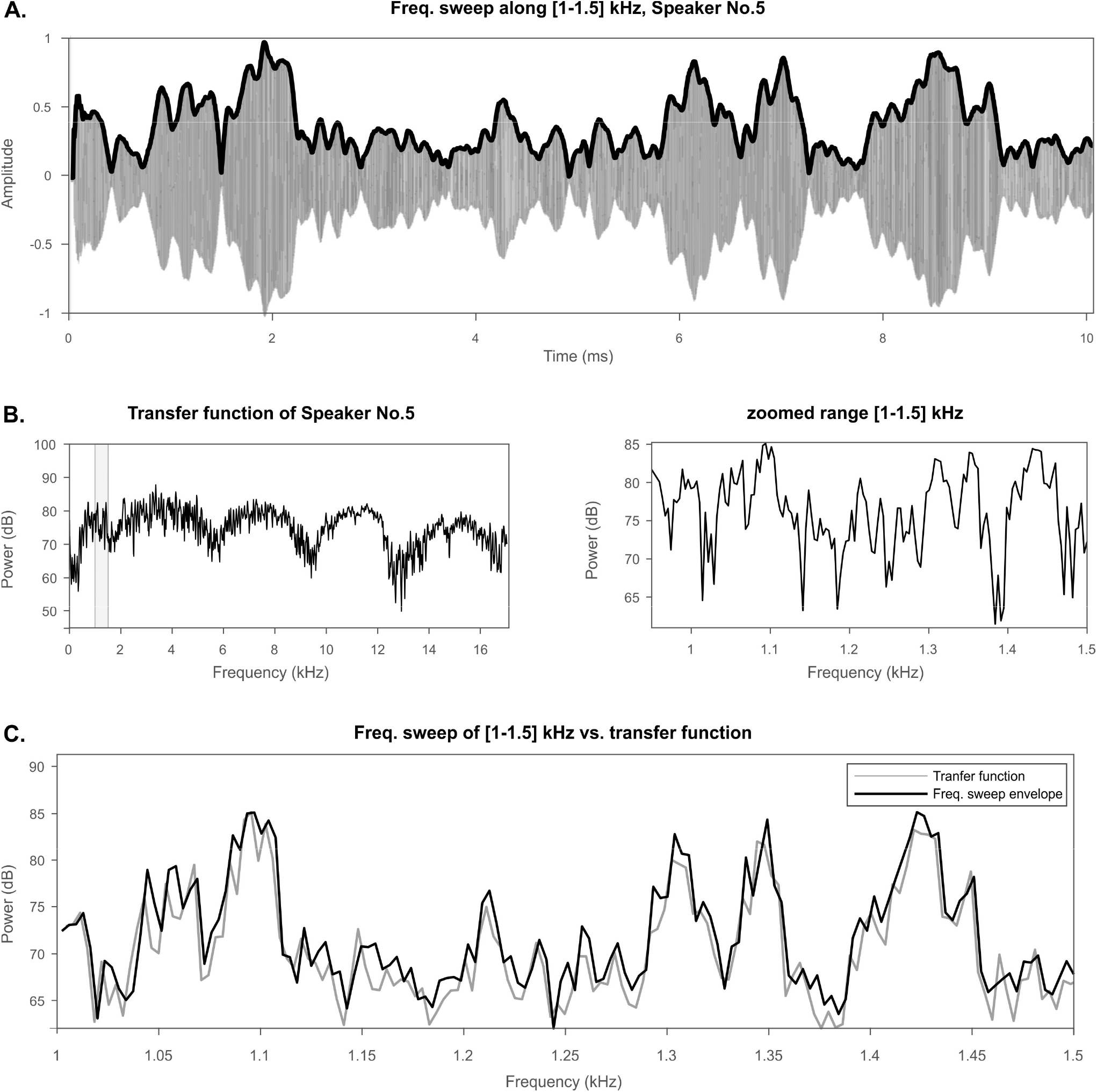
Comparison of the amplitude fluctuations of a frequency sweep with those of the TF measured using the Golay codes. **A**. A Frequency sweep (starting from 1 kHz and ending at 1.5 kHz, duration 10 s) was recorded by a statically located microphone at r=50cm, z=6cm. Bold black line represents the amplitude envelope of the signal **B**. A transfer function was calculated at the same location, using Golay of length 2^14^. The range 1–1.5 kHz (grayed area) is magnified on the right. **C**. The envelope of the sound is compared to the TF, that was converted to dB units using: Power=20·log_10_(Amplitude).

**Supp. fig 10:**
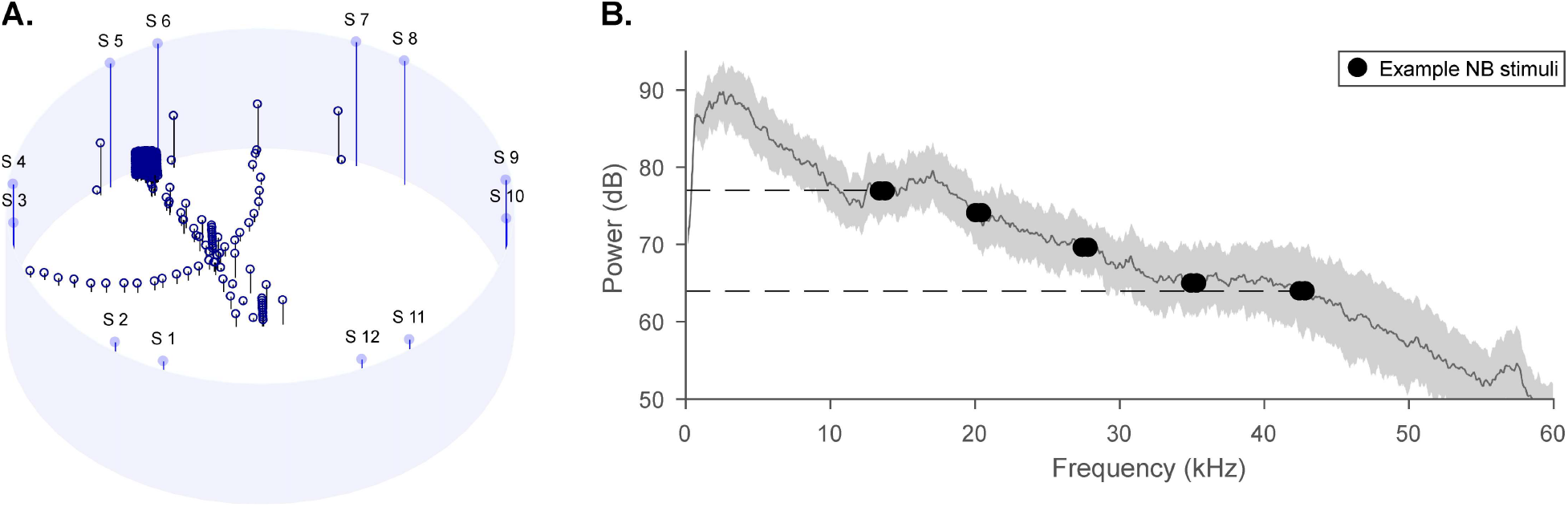
Average TF of 390 location across the arena guides the creation of stable acoustic stimuli. A. Schematic of the 390 recording locations. All 12 speakers were sampled at each location, producing 4680 TFs. **B**. The TFs from A. were averaged (black line is the mean, grey area is the standard deviation). The average TF guides the creation of a stimulus set with minimal power variance (stimuli denoted by black dots). In this example, narrow-band sounds of [12.5 20 27.5 35 42.5] kHz span the frequency range of 30 kHz with power modulation of only 10 dB on average across all the recorded locations.

**Supp. fig 11:**
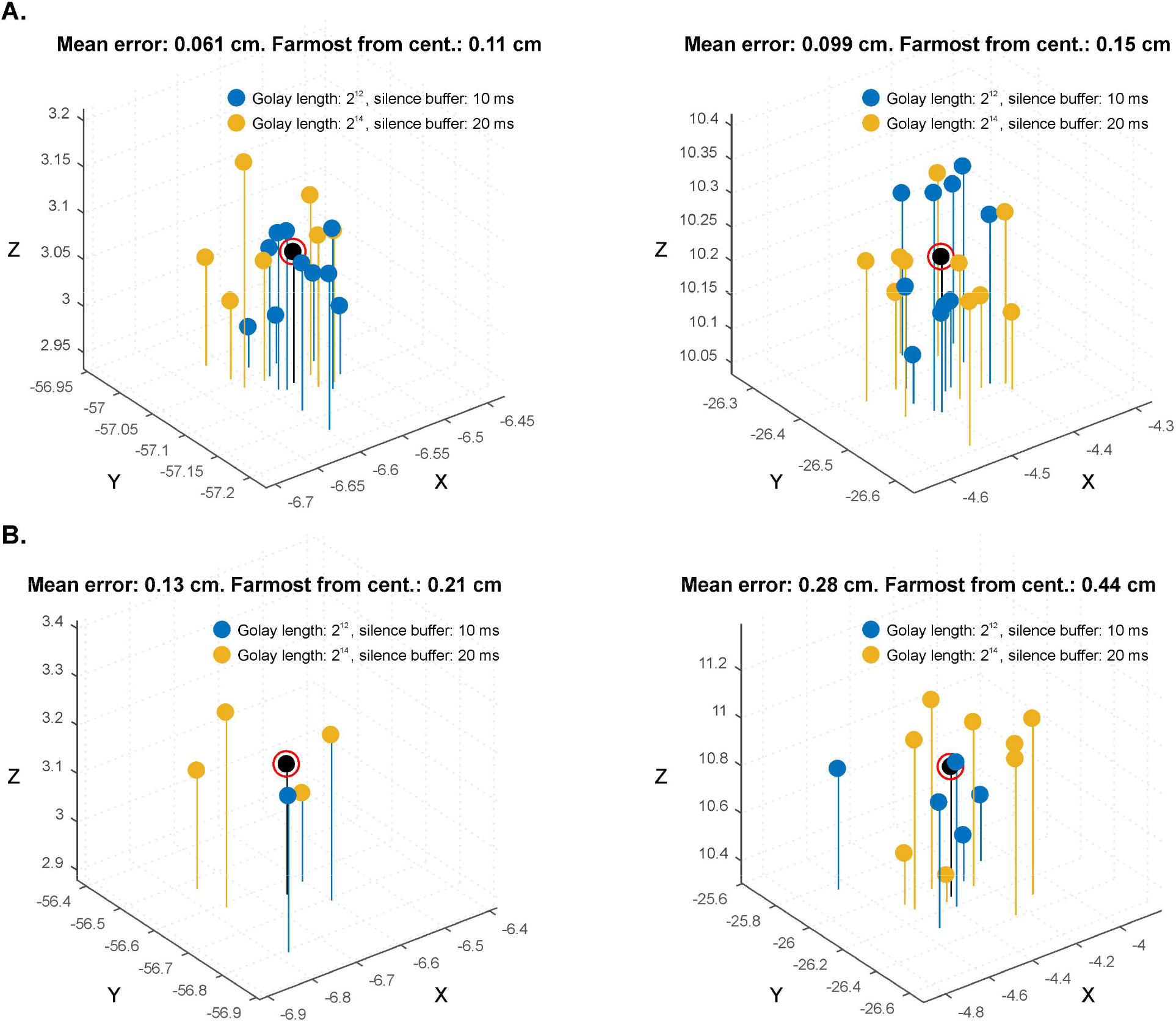
3D Localization of a static microphone achieves sub cm precision, independent on microphone, Golay length or speaker number. A. The 3D localization was tested on a statically placed microphones (B&K in left plots, Headstage in right ones). The localization was obtained based on distances to all 12 speakers. Two different stimulus configurations were tested: Golay of length 2^12^ with silence buffer of 0.1 sec (blue dots) or Golay of length 2^14^ with silence buffer of 0.2 sec (yellow dots). The localization was repeated 10 times for each condition to assess the variance. Axis represent spatial location relative to the center of the arena, in centimeters. B. Same experiment was repeated for localizations based on 3 speakers.

**Supp. fig 12:**
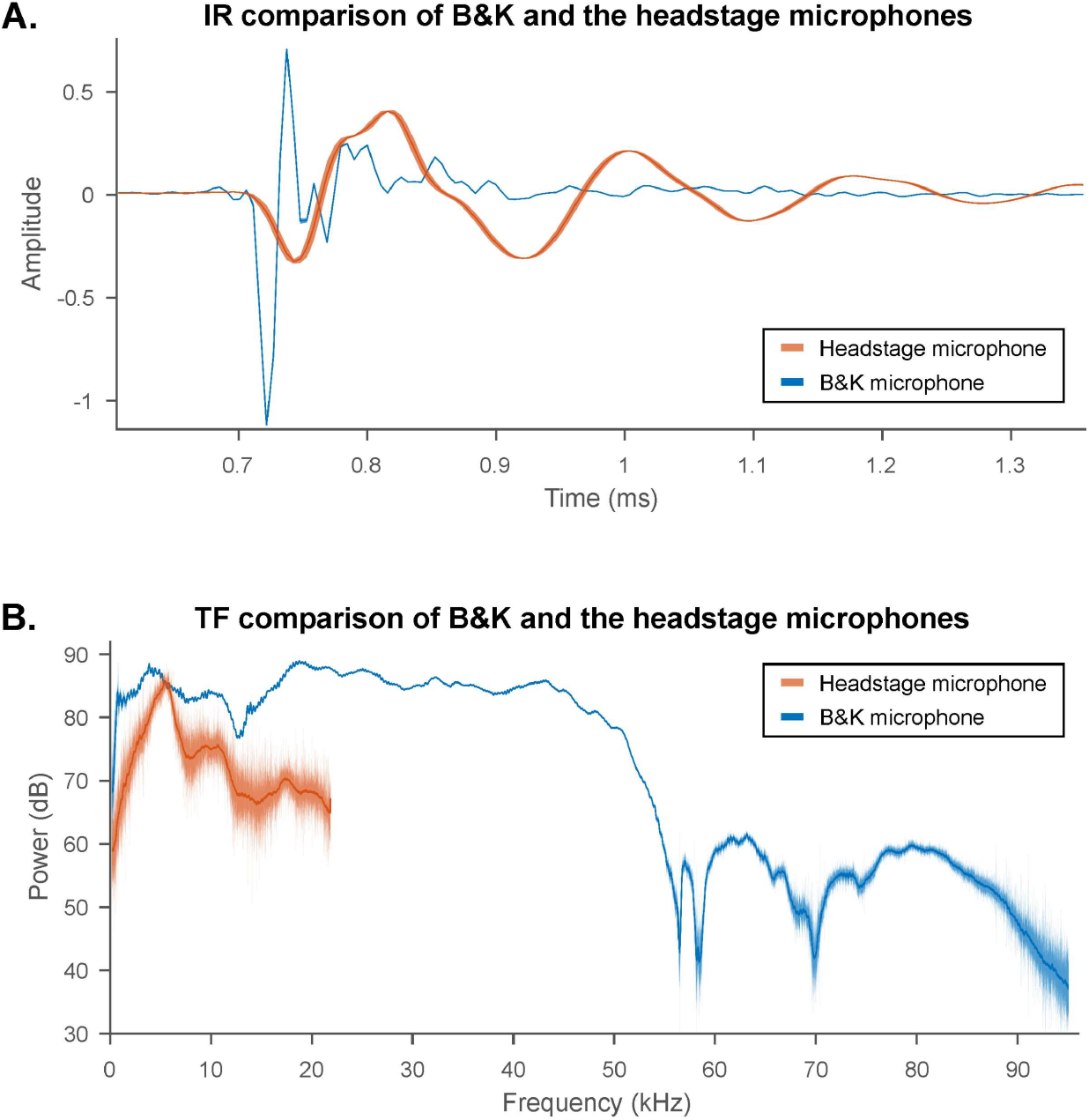
Comparison of the B&K and the HeadStage (HS) microphones TFs. Both microphones were placed 5 cm from the speaker and the IR and TF were measured using Golay of length 2^16^. The measurement was repeated 5 times (solid line indicates the average, the shaded area is standard deviation). Blue color shows the B&K results, orange for HS. **A**. IRs. **B**. TFs.

**Supp. fig 13:**
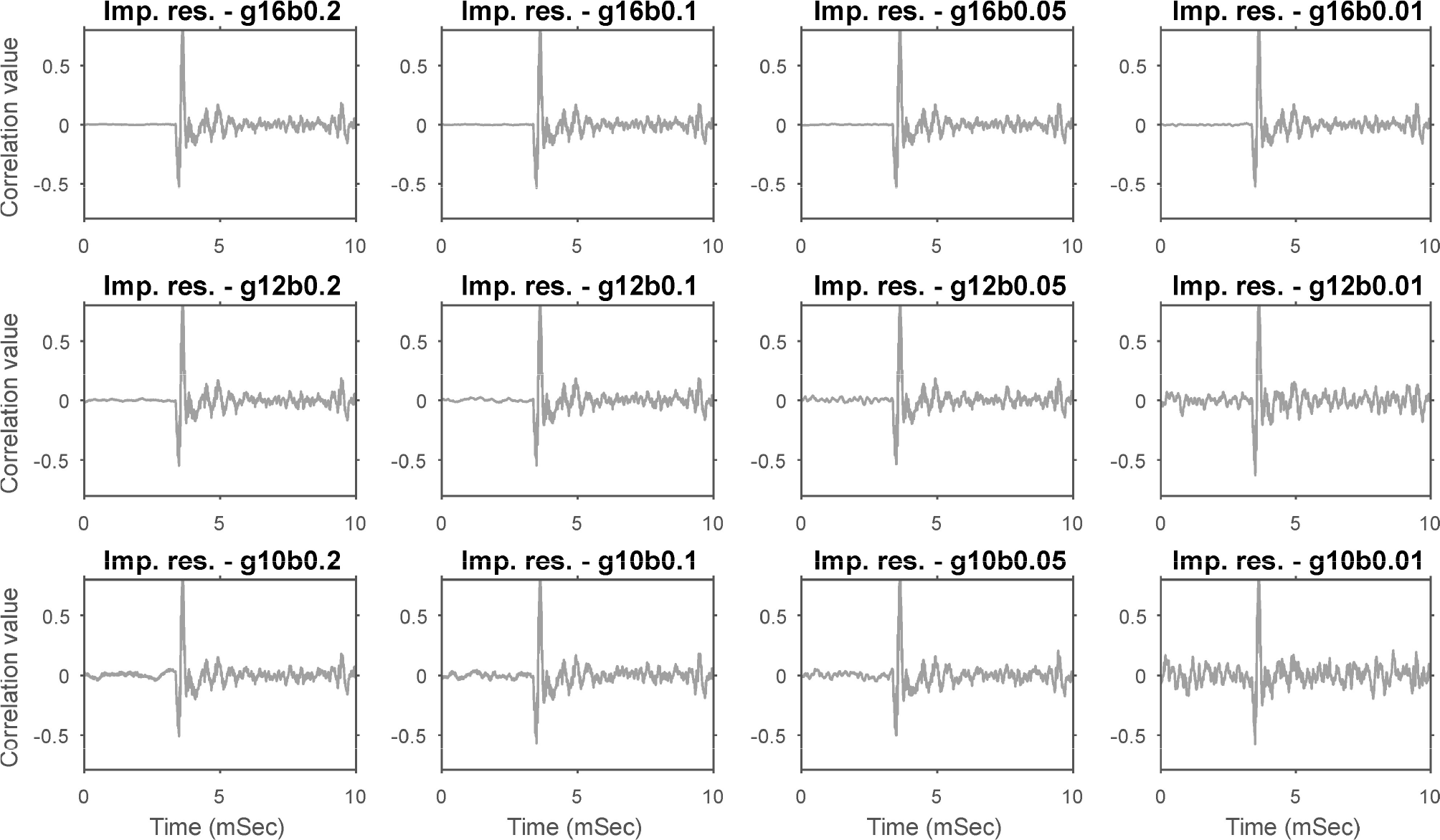
Both Golay and silence buffer length affect the SNR of the impulse response. Impulse response was calculated for Golay cues of varying length (from 2^16^ in upper row down to 2^10^ in the bottom row) and for varying silence buffers (from 200 ms in the left column down to 10 ms in the right column) and rescaled to their individual maximum. The plots represent the first 10 ms of the impulse response, focusing on the direct sound onset at the microphone location. Sounds played and sampled at 192 kHz by B&K microphone.

**Supp. fig 14:**
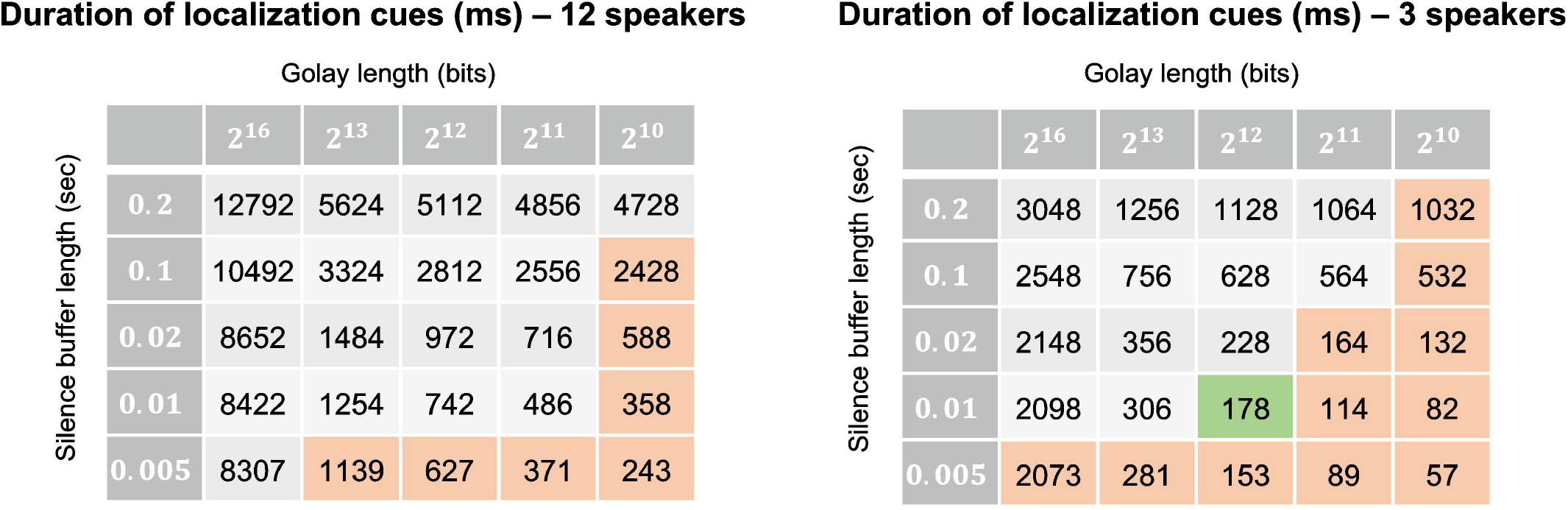
Localization cue length (ms) as a function of Golay length, silence buffer length and number of speakers that were used during the localization. An acoustic localization of the microphone is obtained by playing a localization sound. This sound is comprised of a Golay pairs being played by at least 3 speakers, one at a time, with silence buffers in between. Denote number of speakers by *N*, Golay code length (bits) by *G* and silence buffers (ms) by *t_b_*. The sound duration (in ms) is calculated as follows (sampled as 192 kHz): 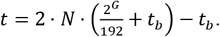 Orange shading stands for localization cues that are too short to successfully localize the microphone (due to low SNR). Green shading is the shortest localization cue that can be used for the localization task.

**Supp. fig 15:**
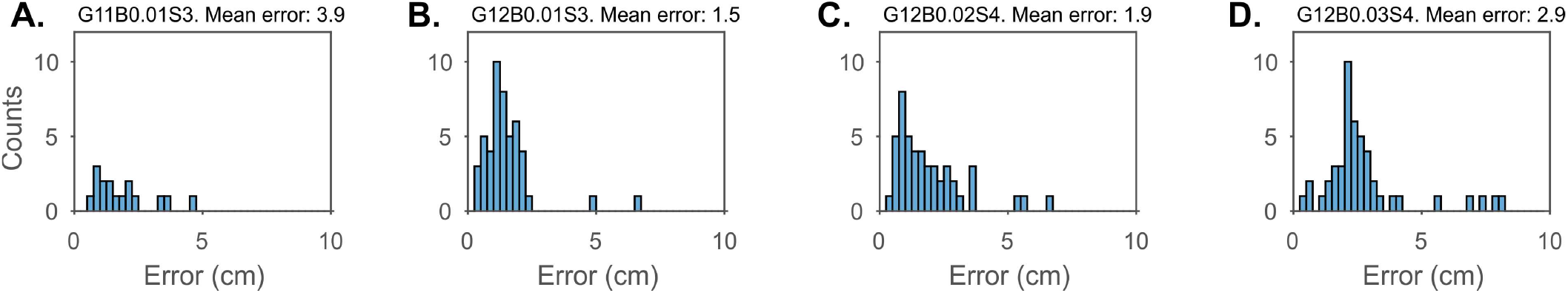
Measured localization error of a freely moving animal as a function of Golay length, buffer length and number of speakers. Four sound configurations were tested for freely moving rat localization. The error was measured as the distance between the acoustical localization and the manual annotation of the microphone location. **A**. Golay of length 2^11^ samples, silence buffer length of 0.01 sec and localization based on 3 speakers, denoted as G11B0.01S3. The test can be viewed in Supp. Movie 1. **B**. Golay = 2^12^, silence buffer = 0.01 sec, 3 speakers. See Supp. movie 2 **C**. Golay = 2^12^, silence buffer = 0.02 sec, 4 speakers. See Supp. movie 3 **D**. Golay = 2^12^, silence buffer = 0.03 sec, 4 speakers. See Supp. movie 4.

